# Long-lasting chromatin and transcriptional states shape the maintenance of memory ensembles

**DOI:** 10.64898/2026.11.16.767121

**Authors:** Ian R. Vogel, Sebastian Krüttner, Natalie J. Guzikowski, Hanqing Liu, Shon A. Koren, Lucy Xu, Stella Sanalidou, Eric C. Griffith, Michael E. Greenberg, Lisa Traunmüller

## Abstract

Memories of recent events are stored within minutes in neuronal ensembles in the hippocampus. Over days to weeks, these memories undergo consolidation, and later retrieval depends on reactivation of a subset of neurons within the ensemble. However, how transient experiences produce lasting molecular and cellular changes in hippocampal ensemble neurons remains unknown. Here, we profile the functional, transcriptional, and gene regulatory states of hippocampal memory ensemble neurons at single-cell resolution across multiple remote timepoints following single-trial contextual fear conditioning. We uncover long-lasting, cell-typespecific chromatin and gene-expression changes after learning that coincide with sustained shifts in synaptic plasticity. Within dentate gyrus granule ensembles, we identify a persistent chromatin– transcriptional program that emerges in a subset of ensemble neurons and is marked by sustained expression of many synaptic genes. This transcriptional signature predicts which neurons will be reactivated during memory recall, suggesting that only a fraction of initial ensemble neurons maintains a molecular state supporting later reactivation. Regulatory elements associated with this persistent state are highly enriched for AP-1 motifs, the cognate binding sites for Fos family transcription factors, even though FOS protein has long since decayed away. Instead, we detect long-lasting FOSB binding at these elements, suggesting a mechanism by which the effects of immediate-early gene–driven transcription are extended beyond the initial induction window. FOSB is particularly enriched in ensemble neurons that maintain enduring gene-expression changes, suggesting it sustains chromatin accessibility and prolonged transcription in neurons reactivatable during memory retrieval. Together, these findings identify molecular mechanisms underlying the maintenance, maturation, and evolution of hippocampal memory ensembles over time.

## INTRODUCTION

Episodic memories are acquired and then consolidated within ensembles of neurons in the hippocampus. Both, loss– and gain-of-function studies have demonstrated that silencing or ablating ensemble neurons disrupts memory recall, whereas artificially reactivating these neurons elicits recall in the absence of a memory retrieval stimulus^1^. While extensive work has revealed how intrinsic neuronal excitability controls recruitment into neuronal ensembles at the moment of learning^1–8^ and has characterized the immediate molecular and physiological changes that accompany memory retrieval^9–14^, the molecular mechanisms by which short-term, labile memories are consolidated into durable memory traces within memory ensemble neurons remain poorly understood.

Activity-dependent neuronal tagging methods have served as indispensable tools in the labeling and manipulation of hippocampal memory ensembles^15^, with approaches based on immediateearly genes (IEGs) such as *Fos* or *Arc* playing a central role in these efforts. These approaches have revealed underlying complexity at the level of the labeled ensembles. For example, while the overlap between populations of neurons that are active during memory encoding with those that are active during memory retrieval exceed chance levels, the overall correspondence between these two populations remains relatively low (typically ∼10-30%)^1,16^. Moreover, reactivation probability varies with the interval between learning and recall, suggesting that the cellular composition and functional architecture of memory ensembles are not static but evolve over the course of days or weeks as a memory of a learned event is refined or consolidated^1,17,18^. Indeed, early post-learning changes in ensemble composition have been linked to the emergence of memory selectivity^19^. Yet, how ensemble identity is stabilized, refined, or selectively maintained over prolonged timescales of days to weeks remained to be elucidated. Moreover, the progressive change of memory representations over time has been shown to be accompanied by structural remodeling, including synapse formation and neuronal circuit reorganization across hippocampal and cortical networks, and reflect changes in memory precision and persistence^16,20^. Thus, defining the molecular mechanisms of hippocampal ensemble dynamics during memory consolidation will be essential for understanding how memories can be maintained over prolonged periods of time.

The activation of IEG-encoded transcription factors (IEG TFs) is a key mechanism by which sensory stimuli regulate neural circuit plasticity. Yet, within the specific neurons that make up memory ensembles, there is an apparent mismatch between the transient nature of IEG mRNA induction and the prolonged duration of ensemble plasticity that can last for weeks to months. As such, this disconnect has contributed to an emphasis on IEGs as markers rather than as mediators of plasticity^1,2,21^. One possibility is that the activity-dependent genes are activated transiently in response to the stimulus and the newly synthesized proteins encoded by these genes are stable and persist within neurons for days to weeks. A second possibility is that by a heretofore uncharacterized mechanism, the activity-dependent gene program is sustained at the transcriptional level and thereby contributes to long lasting memory ensemble function. Still a third possibility is that there are cascades of changing gene expression that give rise to an evolving memory ensemble that encodes, maintains and facilitates memory retrieval over a period of weeks to months. While recent studies have reported long-term transcriptional and epigenomic signatures associated with memory ensembles^11^, how these persistent transcriptional states are achieved and maintained, whether they operate uniformly or heterogeneously within ensemble populations, and if they contribute to ensemble function, remains unclear.

To address these gaps in knowledge, we characterized the functional, transcriptional, and generegulatory landscapes of hippocampal memory ensemble neurons associated with remote memories across time. Using single-nucleus multiome sequencing, we compared ensemble and non-ensemble neurons at one-, two– and ten-days following learning. Importantly, all time points we examined captured molecular features well after the initial IEG and subsequent late-response gene induction. We identified transient as well as surprisingly long-lasting cell-type-specific changes in chromatin accessibility and gene expression. Dentate gyrus (DG) ensemble neurons displayed the strongest and most enduring changes, and these changes paralleled temporally evolving alterations in the functional properties of DG neurons.

Our analysis identified molecular programs of remote phases of memory consolidation during which ensemble plasticity can be maintained in the relative absence of acute stimulus-driven IEG mRNA activation. Strikingly, although many neurons turn on FOS and are engaged during memory encoding, only a subset of these neurons are reactivated upon re-exposure to the same context (10-30%)^1,16^, indicating that memory retrieval depends on a selectively maintained subset of neuronal ensemble cells. Consistent with this, we found that DG memory ensemble cells exhibit significant heterogeneity with respect to maintenance of ensemble-specific molecular and circuit changes. A subset of ensemble neurons maintains persistent expression of the FOSB protein, a distinct paralogue of FOS, over weeks as well as elevated plasticity genes that are likely critical for memory retrieval in a subset of ensemble neurons. The identification of a subset of reactivatable neurons that still exhibit long-lasting transcriptional changes within memory ensembles provides a way forward for further understanding the physiology, synaptic architecture, and molecular features of the retrievable pool of ensemble neurons.

## RESULTS

### Long-term transcriptional profiling of hippocampal fear memory ensembles reveals persistent ensemble-associated gene expression programs

To characterize the gene programs of hippocampal memory ensemble neurons over time, we used an activity-dependent tagging strategy to label learning-activated cells^22^. Using a stereotactic injection protocol we introduced adeno-associated virus (AAV) encoding a Credependent nuclear green fluorescent reporter (AAV-dio-KASH-GFP) into the hippocampus of adult (8-10-week-old) FosTRAP2 mice^22^, enabling 4-hydroxytamoxifen (4-OHT)-dependent labeling of activated neurons within a defined temporal window. Previous studies have shown that this approach efficiently labels the neuronal memory ensemble (or engram) in response to contextual fear conditioning (cFC)^1,2^.

Ten days following injection of the AAV-dio-KASH-GFP virus, the FosTRAP2 mice were subjected to cFC, an established hippocampus-dependent single-trial associative learning paradigm^23^. Hippocampal nuclei were isolated at one-, two– and ten-days post-conditioning. We separately purified NeuN+ learning-tagged GFP+, (hereafter referred to as TRAP+) and untagged nuclei GFP-(hereafter referred to as TRAP-) by fluorescence-activated nuclear sorting (FANS) for single-nucleus multiome sequencing using the 10X Genomics platform (**Fig. 1a, S1a**). Samples were obtained from three independent cohorts for each timepoint to generate paired transcriptome (snRNA) and chromatin accessibility (snATAC) profiles. One of the samples at twodays post cFC did not meet snRNA-seq quality control metrics and was excluded from further differential gene expression analysis. Overall, we obtained data from a total of 170,450 nuclei that passed quality control metrics, with transcriptional and chromatin profiles highly reproducible across replicates (**Fig. 1b, S1b-e, Supplementary Table 1**). This large number of neuronal nuclei characterized at single cell-resolution enabled comprehensive cell-type-specific analysis of differential gene expression and identification of gene regulatory elements in memory ensemble neurons.

**Figure 1:**
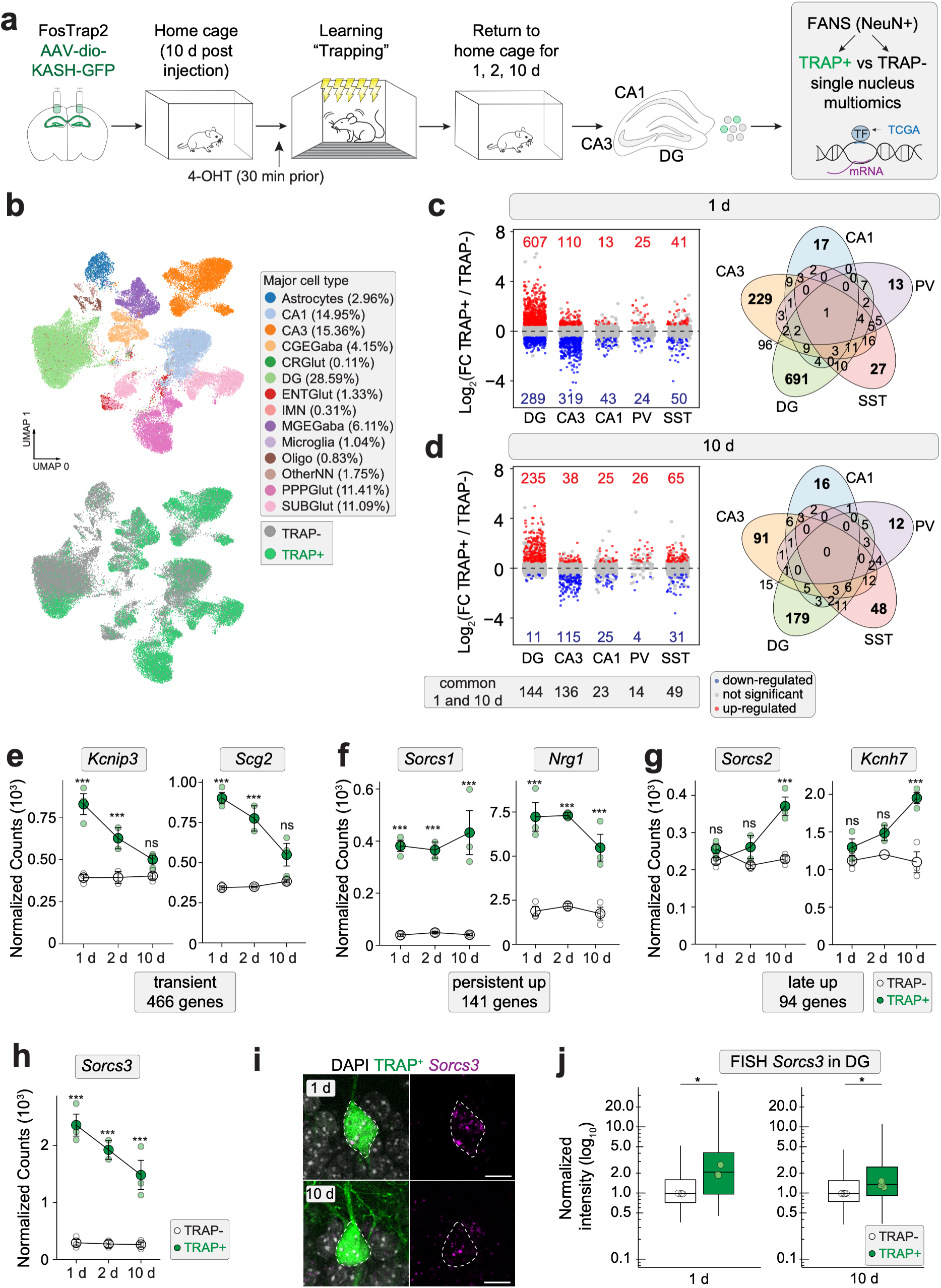
Dynamic cell-type-specific transcriptional remodeling in hippocampal memory ensembles. a) Schematic of experimental strategy for activity-dependent labeling and time course of single-nucleus multiomic profiling of hippocampal fear memory ensembles using FosTRAP2 mice at one, two or ten days following contextual fear conditioning. Upon sample collection, fluorescence-activated nuclear sorting (FANS) was used to sort NeuN+ nuclei into either TRAP+ (GFP+) or TRAP-(GFP-) populations. b) UMAPs of integrated snRNA datasets display major cell classes detected and their respective TRAP status. Percentage of total nuclei analyzed are indicated. c) (Left) Jitter plots of pseudobulk DESeq2 differential gene expression analysis between TRAP+ and TRAP-neurons within the five major cell classes investigated at one day. Genes significantly up– or down-regulated in TRAP+ neurons are marked in red or blue, respectively (Log_2_FC ± 0.5, adjusted p-value ≤ 0.01, two-sided Wald test with Benjamini Hochberg correction for multiple testing). (Right) Venn diagrams showing the number of TRAP+ differentially expressed genes that are commonly shared or unique to each of the five cell types analyzed one day post cFC. d) As c) but for ten days post cFC. Numbers of differential genes commonly affected at one and ten days for each cell class are shown below. e-h)DESeq2 normalized counts of example genes exhibiting transient (Kcnip3, *Scg2*), persistent (*Sorcs1, Sorcs3, Nrg1*) or late (*Sorcs2, Kcnh7*) up-regulation in DG neurons. Signal from TRAP+ (green) and TRAP-neurons (white) is shown. Larger circles display means across replicates, whereas smaller circles represent individual animals. Error bars depict ± SEM. Asterix indicate significance *** < 0.001, ns = non-significant, two-sided Wald test with Benjamini Hochberg correction for multiple testing i) Representative fluorescent in-situ hybridization images showing *Sorcs3* mRNA in TRAP+ DG neurons at one or ten days post cFC. Scale bar is 10 µm. Quantification of normalized *Sorcs3* intensity values in TRAP+ and TRAP-DG neurons at one or ten days post cFC, Box plots depict median intensity of all cells, circles indicate individual animals, One day: N = 3 animals, n = TRAP+: 502 cells / TRAP-: 11444 cells; Ten days: N =4 animals, n = TRAP+: 757 cells / TRAP-: 10486 cells. Significance assessed with paired t-tests comparing TRAP+ vs TRAP− groups. 1 day, p =0.0486; 10 days, p =0.0108.

Graph-based clustering resolved major neuronal cell classes that were similar to those detected in prior hippocampal single-cell studies^24,25^. For subsequent analyses, we focused on five major neuronal cell types robustly represented in TRAP+ and TRAP-populations across all replicates: DG granule cells, cornu-ammonis (CA) 3 and CA1 excitatory neurons, as well as somatostatin (SST)-and parvalbumin (PV)-positive interneurons. Differential pseudobulk gene expression comparison between TRAP+ and TRAP-neurons within each cell class revealed extensive gene expression changes at one-day post-cFC (**Fig 1c, left panel),** (Log_2_FoldChange ± 0.5 and Padj <0.01, **Supplementary Tables 1-2,**). These differentially expressed genes (DEGs) were highly cell-type-specific, with the largest number observed in DG granule cells and the remaining changes largely confined to individual cell types (**Fig. 1c, right panel**). Gene Ontology analysis revealed that many of these DEGs encode synaptic proteins, potassium channels, and secreted factors (**Fig. S2a**), suggesting possible mechanisms of memory encoding within the memory ensemble neurons.

Surprisingly, comparison of the TRAP+ and TRAP-neurons at two– and ten-days following training revealed memory ensemble-associated gene expression patterns similar to those observed after one day, with DG granule cells displaying the most pronounced changes in gene expression (**Fig. 1d, Fig. S2b-c, Supplementary Tables 1-2,**). While the number of DEGs declined at later timepoints, >50% of DEGs identified at ten days also showed significant expression differences between TRAP+ and TRAP-neurons at one day (**Fig. 1d**). The magnitude of differential gene expression in the DG was much greater than that detected in CA1 and CA3. This allowed us to classify upregulated genes in DG ensemble neurons into three groups based on their overall expression kinetics as: transient – detected as differentially up-regulated at one day (466 genes, e.g. *Kcnip3*, *Scg2*, **Fig. 1e**), persistent – significantly up-regulated at one and ten days (141 genes, e.g. *Sorcs1, Nrg1*, **Fig. 1f**), or late – significantly up-regulated only at ten days (94 genes, e.g. *Sorcs2, Kcnh7*) (**Fig. 1g**). Interestingly, parts of the late program mainly emerged from a small subcluster of DG neurons (**Fig. S2d**) and was also associated with genes implicated in synaptic function. Given the striking effects on persistently expressed genes over time, we independently validated the persistent gene expression program in TRAP+ neurons by fluorescence in-situ hybridization (FISH). We focused our attention on *Sorcs3*, as this was our most differentially expressed, persistent gene at ten days in DG. We observed an enrichment for *Sorcs3* mRNA in TRAP+ neurons at both one and ten days after cFC (**Fig. 1h-j**). Taken together, these findings suggest that memory ensemble neurons undergo extensive and temporally structured transcriptional remodeling well beyond the initial IEG response, consistent with sustained molecular programs that may support long-term synaptic plasticity.

A comprehensive overview of cell-class-specific transcriptional changes across all neuronal populations is provided as a supplementary document, offering a rich resource for further exploration of ensemble-associated gene regulation over time (**Supplemental Resource document**). Given the magnitude of persistent and late-emerging ensemble-associated gene expression changes in DG granule cells, we focused our subsequent analysis on this population. The substantial number of genes with functions relevant to synaptic plasticity in the DG, at times as long as ten days after learning (**Fig. S2a-b**), suggests that one trial associative learning leads to long-lasting transcriptional responses that likely contribute to the re-wiring of the neuronal circuitry and alterations in synaptic connectivity and intrinsic properties of neuronal ensemble cells during memory consolidation.

### Persistent functional remodeling of DG fear ensembles

We next investigated whether the temporally evolving memory ensemble-associated gene expression program is accompanied by durable changes in neuronal circuit properties during the early and late phases of memory consolidation in the DG. To this end, we adapted our experimental paradigm to virally label TRAP+ cells in FosTRAP2 mice using a Cre-dependent nuclear blue fluorescent protein (AAV-dio-nBFP). We then performed whole-cell patch-clamp recordings from neighboring TRAP+ and TRAP-DG cells in acute hippocampal slices at two– and ten-days post-training, allowing for sufficient accumulation of the fluorophore in the nucleus for the identification of ensemble neurons (**Fig. 2a**). At two days post learning, TRAP+ neurons exhibited significant increases in the frequency of miniature excitatory and inhibitory post-synaptic currents (mEPSCs and mIPSCs, respectively) relative to TRAP-neurons, with minimal differences in event amplitude (**Fig. 2b, d, S3a-d**). In addition, current-clamp recordings revealed reduced intrinsic excitability of TRAP+ cells compared to TRAP-control neurons (**Fig. 2f, S3e, g, i**). Thus, two days after cFC, DG ensemble neurons exhibit increased synaptic connectivity but reduced intrinsic excitability, reflecting coordinated synaptic adjustments that may stabilize ensemble function. This decreased excitability may limit the incorporation of DG ensemble neurons into additional memory traces during early consolidation.

**Figure 2:**
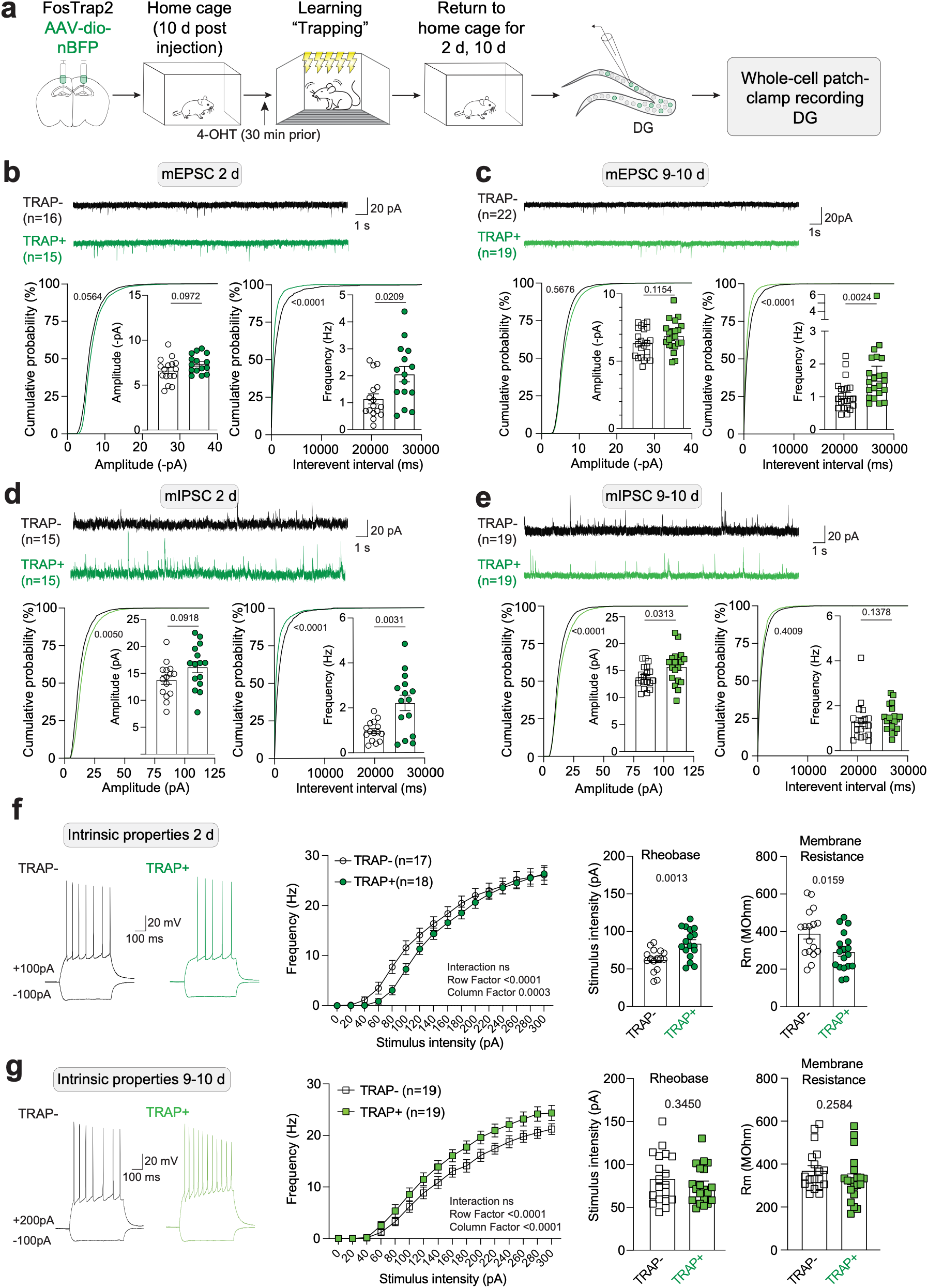
Evolving and persistent and electrophysiological properties in dentate gyrus memory ensembles. a) Schematic of experimental design for whole-cell patch-clamp recording from TRAP+ and neighboring TRAP-DG granule cells at two or nine/ten days following cFC. b-c) Representative traces of TRAP+ (green) and TRAP-(black) DG neurons. Cumulative distributions and mean amplitude and frequency of miniature excitatory postsynaptic currents (mEPSC) at two (b) or nine/ten days (c) following cFC. Two days: N=4 animals; TRAP+ n=15 cells, TRAP-n=16 cells, Nine/ten days: N=6 animals; TRAP+ n=19 cells, TRAP-n=22 cells. d-e) Representative traces of TRAP+ (green) and TRAP-(black) DG neurons. Cumulative distributions and mean amplitude and frequency of miniature inhibitory postsynaptic currents (mIPSC) at two (d) or nine/ten days (e) following cFC. Two days: N=4 animals; TRAP+ n=15 cells, TRAP-n=15 cells, Nine/ten days: N=6 animals; TRAP+ n=19 cells, TRAP-n=19 cells. f-g) Representative firing patterns of DG granule cells in response to single 500 ms long hyperpolarizing and depolarizing injected currents. Intrinsic property measurements at two (f) and nine/ten days (g) post cFC. Frequency in Hz of action potential firing in response to increasing current injections (statistics of Two-way Analysis of variance (ANOVA) are indicated). Rheobase and membrane resistance are displayed for TRAP+ and TRAP-neurons. Two days: N=5; TRAP+ n=18, TRAP-n=17, Nine/ten days: N=7; TRAP+ n=19, TRAP-n=19. Unless otherwise indicated, statistics for each comparison were calculated based on the appropriate unpaired t-test based on normality distribution and standard deviation, KolmogorovSmirnov test for cumulative distributions. Mean ± SEM are shown.

Electrophysiological recordings at nine/ten days post cFC revealed a continued evolution of DG ensemble properties. While the elevated mEPSC frequency observed in TRAP+ DG ensemble neurons at two days persisted at ten days, mIPSC frequency returned to baseline levels (**Fig. 2c, e**). At ten days, we detected a moderate but significant increase in mIPSC amplitudes with faster rise times (**Fig. 2e, S3c**), suggesting alterations in GABA-A receptor subunit composition or localization. Current-clamp recordings also revealed a further shift in intrinsic excitability over time: whereas TRAP+ DG ensemble neurons exhibited reduced excitability at two days, they displayed increased excitability relative to TRAP-neurons by ten days post-learning (**Fig. 2g, S3f, h, i**). These prolonged synaptic changes may contribute to the preservation of input-specific reactivation dynamics, while the higher intrinsic excitability at ten days may enable ensemble reactivation in response to incomplete cues or support a capacity for allocation to additional ensembles.

Intriguingly, we identified gene expression patterns whose kinetics closely follow the observed changes in functional properties of the DG granule cell ensemble neurons. These include persistent genes associated with regulating excitatory synaptic transmission via AMPAR clustering (e.g. *Sorcs1*, *Sorcs3*, *Nptx2*^26–28^), transient genes involved in the regulation of inhibition onto excitatory neurons (e.g. *Scg2*, *Gabrd*^29,30^) or genes that regulate neuronal excitability (e.g. *Kcnip3*, *Kcnj2*^31–33^). Taken together, our transcriptional and functional characterization of the hippocampal memory ensemble reveals previously unrecognized and distinct long-lasting changes in gene expression that evolve alongside significant alterations in synaptic plasticity.

### Persistent engram-associated accessible chromatin regions are enriched for AP-1 binding motifs

To further investigate the mechanisms that mediate persistent ensemble-associated gene expression, we leveraged our single-nucleus multiome dataset to analyze chromatin accessibility across cell types at one, two-, and ten-days post-cFC. Intra-cell-type comparison of TRAP+ and TRAP-neurons at each timepoint identified thousands of cell-class-specific differentially accessible regions (DARs) in DG, CA3, CA1, and MGE-GABA (which combines SST and PV neurons) (**Fig. S4a, Supplemental Resource Document**). Mirroring our gene expression findings, DG granule cells exhibited particularly extensive and long-lasting changes in chromatin accessibility, with pseudobulked differential accessibility analysis identifying ∼18,000 highconfidence DARs with increased accessibility in TRAP+ DG neurons (**Fig. 3a-b**). These ensemble-accessible regions exhibited distinct temporal kinetics, with 6,711 transient regions (differentially accessible exclusively at one day) and 8,553 semi-transient peaks (differentially accessible at both one and two days). Remarkably, we also identified 2,031 persistent peaks that continue to be differentially accessible from day one to day ten. Despite the variable kinetics with which these putative regulatory regions open and close, there were very few newly emerging accessible regions at later times (**Fig. 3a, S4a**).

**Figure 3:**
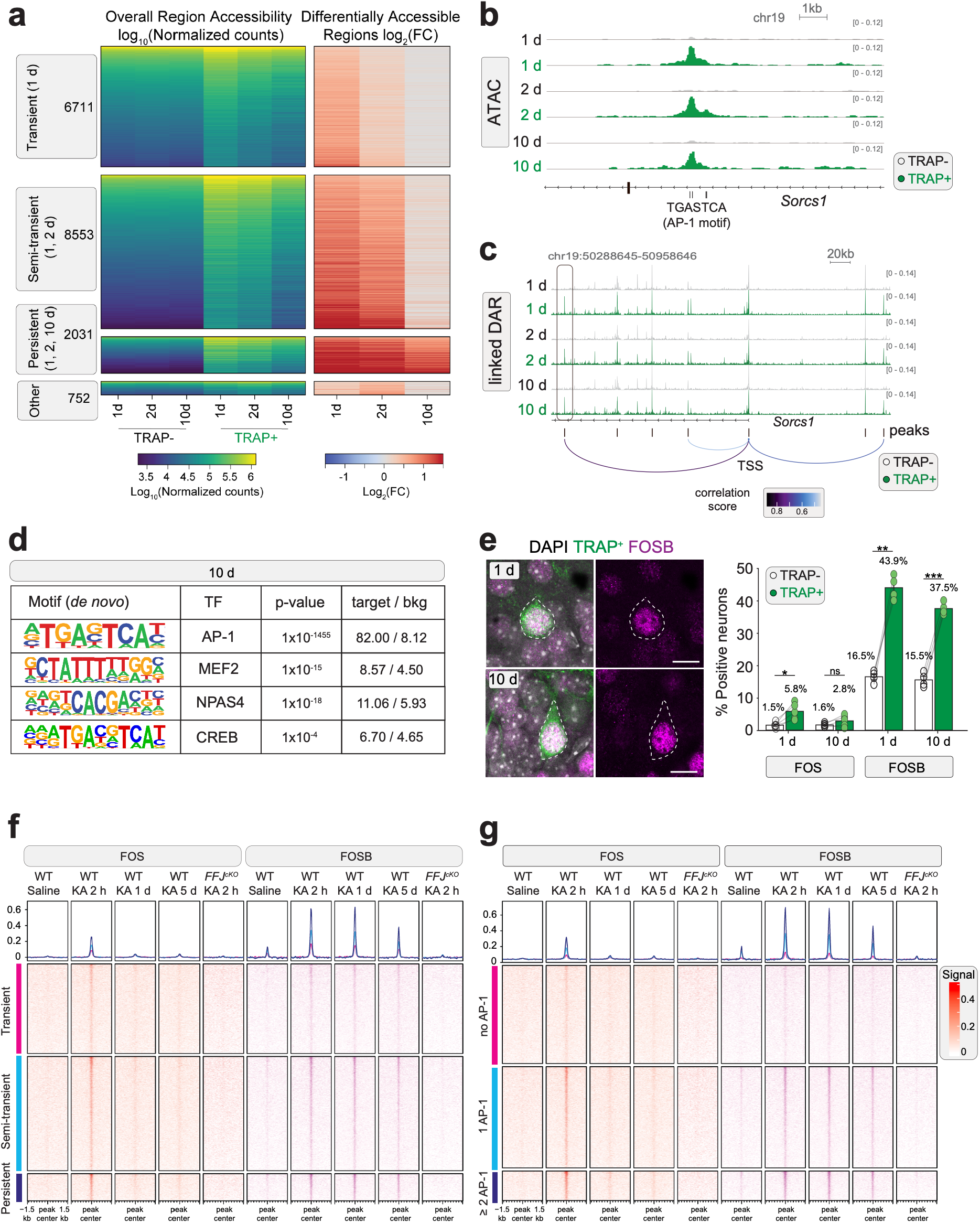
Persistent memory ensemble chromatin states are associated with sustained FOSB binding. a) Heatmaps of pseudobulked differential chromatin accessibility analysis, log_10_(DESeq2normalized counts) (left) and TRAP-enriched log_2_(Fold Change) (right) across excitatory DG neurons with distinct transient, semi-transient, persistent and other accessibility kinetics (DEseq2 adjusted p-value ≤ 0.05, two-sided Wald test with Benjamini Hochberg correction for multiple testing). b) Genome browser tracks of chromatin accessibility in the intronic region of the persistent *Sorcs1* gene in TRAP+ and TRAP-neurons across time. Black vertical lines depict AP-1 motif positions. c) ArchR generated ATAC-seq tracks and peak-to-gene linkages for significant Pearson correlation of three TRAP-enriched peaks upstream or within the *Sorcs1* gene in DG neurons. TSS = Transcription Start Site. Black vertical lines = all peaks detected. Colored arcs depict TRAP+ Enriched DARs correlated with Sorcs1 expression. The peak highlighted in 3b) is indicated by the left box. d) De novo motif analysis using HOMER for the top enriched motifs at ten-day post cFC. Motifs, associated transcription factors (TF), p-values and percentage of enrichment over background (bkg) are presented. Number of peaks at ten days: 2,089. e) (Left) Representative image of labeled TRAP+ dentate gyrus neurons (green), FOSB protein staining (magenta) and DAPI (grey) at one or ten days. Scale bar = 10 µm. (Right) Quantification of FOS and FOSB protein positivity rate in TRAP+ and TRAP-dentate gyrus granule cells. FOS one day: N= 5 animals; n = TRAP+: 829 cells / TRAP-: 8671 cells, FOS ten days: N = 5 animals; n = TRAP+: 800 cells /TRAP-:7018 cells. FOSB one day: N = 4 animals; n = TRAP+: 215 cells / TRAP-: 2963 cells, FOSB ten days: N = 4 animals; n = TRAP+: 369 cells / TRAP-: 3196 cells. Significance assessed using twoway mixed-effects ANOVA performed separately for each protein, TRAP status as withinanimal factor and timepoint as between-animal factor, with post hoc paired t-tests with Benjamini–Hochberg correction. %Fos⁺ cells (TRAP⁺ vs. TRAP⁻), Padj = 0.011 (1 day) and 0.14 (10 days). %FosB⁺ cells, Padj = 0.0002 (1 day) and 1.97 × 10⁻⁶ (10 days). A significant effect of time was observed only for FosB in TRAP⁺ cells (1 day vs. 10 days, adjusted p = 0.0208). f) CUT&RUN profiling of FOS or FOSB binding following saline (2 h) or kainic acid-induced neuronal activation (2 h, 1 d, 5 d) in WT animals or FFJ conditional knock-out mice (2 h). Binding of FOS or FOSB at either transient (pink), semi-transient (light-blue) or persistent (dark-blue) open chromatin regions identified after cFC is displayed. Top: aggregation of binding over all peaks, bottom: accessibility at individual peaks around peak center. n = 2 mice per biological replicate, Four biological replicates per WT, and two biological replicates for FFJ^cKO^. g) CUT&RUN profiling of FOS or FOSB binding as in f) but at peaks defined by the number of AP-1 motifs: pink no AP-1, light-blue one AP-1, and dark-blue more than two AP-1 motifs.

Across all timepoints, DG ensemble-enriched DARs were predominantly intronic or intergenic, with approximately one-third located within 500 kb of a DEG locus, suggesting that many of these open regions are distal regulatory elements that control gene expression (**Fig. S4b**). DAR–gene correlation analysis using ArchR^34^ supported this relationship, linking 1,513 unique DARs with 377 unique up-regulated DEGs (DEG-linked-DARs, r>0.45, Padj <10^-23^). Strikingly, over 50% of all upregulated DEGs were significantly correlated with at least one nearby DAR in TRAP+ DG ensemble neurons (**Fig. 3c**, **Supplementary Table 3**). Taken together, our findings indicate that in response to cFC, hippocampal ensemble neurons undergo an increase in chromatin accessibility at thousands of regions across the genome. Surprisingly, many of these increases are long-lasting and likely give rise to sustained programs of gene expression that contribute to memory consolidation.

To gain insight into the regulatory mechanisms by which cFC promotes chromatin opening in TRAP+ neurons, we performed *de novo* motif analysis^35^ on genomic regions exhibiting enhanced accessibility in the TRAP+ neurons. Surprisingly, across all time points, even ten days following cFC, these sites showed a striking enrichment for the AP-1 motif relative to matched nonensemble-selective sites, whereas enrichment observed for other activity-regulated TFs was substantially weaker (for DG at ten days post cFC: FOS ∼10.1-fold, followed by MEF2 ∼1.9-fold, and NPAS4 ∼1.8-fold, CREB ∼1.4-fold; **Fig. 3d, Fig. S4c**). Moreover, persistent DG ensembleaccessible sites were strongly enriched for the AP-1 motif relative to transient sites (i.e. 87.9% of persistent sites harbored at least one AP-1 motif versus 36.6% of transient sites) and were roughly sevenfold more likely to harbor one or multiple AP-1 motifs (**Fig. S4d**). No additional sequence motifs were found to distinguish persistent from transient regions, suggesting that prolonged accessibility is primarily associated with AP-1 motif density. Together, these findings raised the possibility that AP-1 family members might play an ongoing role in mediating the long-lasting chromatin accessibility and gene expression programs observed in hippocampal memory ensembles.

### Persistent FOSB expression in DG ensemble neurons

Given the strong enrichment of AP-1 motifs in ensemble-associated regulatory regions, we next asked whether the AP-1 motif binding factors are required for contextual fear memory consolidation. To this end, we first employed triple-conditional-FOS, FOSB, JUNB KO mice (*Fos^fl/fl^;Fosb^fl/fl^;Junb^fl/fl^*, hereafter FFJ)^29^ to achieve the deletion of these strongly inducible AP-1 family members in a spatiotemporally controlled manner. The hippocampi of adult FFJ mice were bilaterally injected with AAV expressing Cre (FFJ-KO) or a catalytically inactive ΔCre (FFJ-WT). After a ten-day recovery period, mice were subjected to fear conditioning, and AP-1 family member loss was found to significantly reduce memory recall one day post-conditioning (**Fig. S5a, b**). Thus, consistent with prior reports^36,37^, we conclude that AP-1 family members are essential for the encoding and or recall of fear memories in the hippocampus.

While these findings suggest that *Fos* or a related family member mediates the induction of gene expression in TRAP+ neurons in response to cFC, they raise the question of how the expression of these genes is maintained in TRAP+ neurons for many days. This is particularly intriguing as FOS protein is expressed shortly after cFC and employed for the labeling of the TRAP+ population, but is expected to have decayed away within hours post cFC. One possibility is that in DG neurons, in response to cFC, the FOS protein is induced and stable or recurrently reinduced to sustain gene regulation. In this case, the FOS protein should be detectable in TRAP+ neurons at one or ten days after cFC. A second possibility is that FOS is necessary for an initial priming event that licenses regulatory regions for activation, but that other long-lived AP-1 family members, or unrelated TFs, mediate gene activation during memory consolidation.

To investigate if sustained or recurrent FOS induction occurs at remote times following cFC, we immunostained hippocampal sections for FOS at one– and ten-days post-cFC and quantified FOS expression in TRAP+DG granule cells. At both time points, only a small fraction of TRAP+ ensemble neurons express FOS (5.8% at one day, 2.8% at ten days; **Fig. 3e**). While this represents a weak but statistically significant enrichment over TRAP-cells at one day (1.5%, Padj = 0.011) it does not at ten days (1.6%, Padj = 0.14). By contrast, immunostaining for FOSB, a related Fos family TF, revealed a striking difference: 43.9% of TRAP+ neurons are FOSB-positive at one day, and 37.5% even ten days post cFC, significantly exceeding the level of FOSB in neighboring TRAP-neurons (16.5%, Padj = .002 and 15.5%, Padj = 1.9×10^-6^; **Fig. 3e**). Notably, ΔFOSB, a truncated *FosB* splice variant, has been found to be stable at the protein level, with a reported in vivo half-life of approximately eight days^38,39^, providing a potential explanation for the sustained FOSB expression.

We next sought to determine if the AP-1 motif-containing open regulatory regions in TRAP+ neurons bind FOS and/or FOSB at various times after cFC. However, the low number of memory ensemble neurons (∼1-2% of total hippocampal neurons) precluded direct assessment of TF occupancy in ensemble neurons using methods such as CUT&RUN. Therefore, as an alternative approach, we exposed mice to kainic acid to obtain maximal, synchronous induction of the activity-dependent gene program and performed CUT&RUN^40^ for FOS and its close relative FOSB on isolated NeuN+ DG nuclei at two hours, one day, or five days following the injection of kainic acid. We further assessed FOS and FOSB binding at the DARs we previously identified in our DG TRAP+ neurons. Within two hours of kainic acid treatment, we observed strong inducible binding of FOS at the sites that we found to be selectively open in TRAP+ neurons, with the highest binding of FOS at the subset of these sites that exhibit persistent opening and/or multiple AP-1 motifs (**Fig. 3f-g**, FOS). However, consistent with the documented kinetics of FOS protein synthesis and degradation^41^, FOS binding at these sites reverts to baseline by one– and five-days post-seizure (**Fig. 3f-g**, FOS). By contrast, FOSB binding persists at these same sites at the later time points. Notably, the magnitude and durability of FOSB binding scales with the ensemble specific persistence of region accessibility and density of FOS/AP-1 motifs within these regulatory sites (**Fig. 3f-g**, FOSB). Importantly, two hours following kainic acid treatment we detected no signal at these sites when the CUT&RUN analysis was performed using nuclei from FFJ mice injected with Cre or in the IgG control, indicating that the binding of FOS and FOSB to these sites is specific (**Fig. 3f-g, Fig. S5d**). These findings, taken together with our functional and immunohistochemical data, suggest that while FOS binding immediately after cFC leads to the opening of AP-1 motif-containing regulatory elements in many TRAP+ DG neurons, FOSB binding then sustains the opening of these elements long after the initial cFC stimulus in a subpopulation of TRAP+ neurons, thereby maintaining plasticity-associated gene expression for prolonged periods of time.

### Heterogeneous decay of DG fear ensemble chromatin state

Analysis of FOSB protein expression at one– or ten-days post cFC revealed that ∼40% of DG ensemble neurons retain FOSB expression (**Fig. 3e**). Interestingly, we detected a significant range in the number of cells as well as their intensity of the FOSB signal within the DG memory ensemble neurons. We reasoned that the heterogeneity in FOSB abundance and its binding at persistent DARs might give rise to heterogeneity in the chromatin and gene expression states within ensemble neurons, thus influencing ensemble durability. To address this possibility, we used the ChromVAR^42^ package to assess the accessibility of AP-1 motifs within individual DG neurons to generate per-cell ChromVAR “scores” that reflect the overall accessibility of AP-1associated regulatory elements in a given cell. Across all time points, TRAP+ DG neurons exhibit significantly higher AP-1 ChromVAR scores than TRAP-neurons, and the range in ChromVAR scores in the TRAP+ neurons is markedly broader, indicating that within the memory ensemble neurons, there is substantial intra-ensemble heterogeneity with respect to the degree of AP-1 site opening (**Fig 4a, Fig. S5e**). While this intra-ensemble heterogeneity is evident as early as one day after cFC, the pattern of chromVAR scores appears to be redistributed over time. Interestingly, although the fraction of TRAP+ DG neurons with high AP-1 ChromVAR scores decreases over time, a significant portion (∼25%) of TRAP+ neurons at ten days have scores similar to those observed one day after cFC (**Fig. 4a, Supplementary Table 3**). Thus, despite a population-level reduction in ensemble-associated chromatin accessibility over time (**Fig. 3a**), the accessibility of AP-1 regulatory sites is selectively maintained at a high level in a significant subset of ensemble neurons.

**Figure 4:**
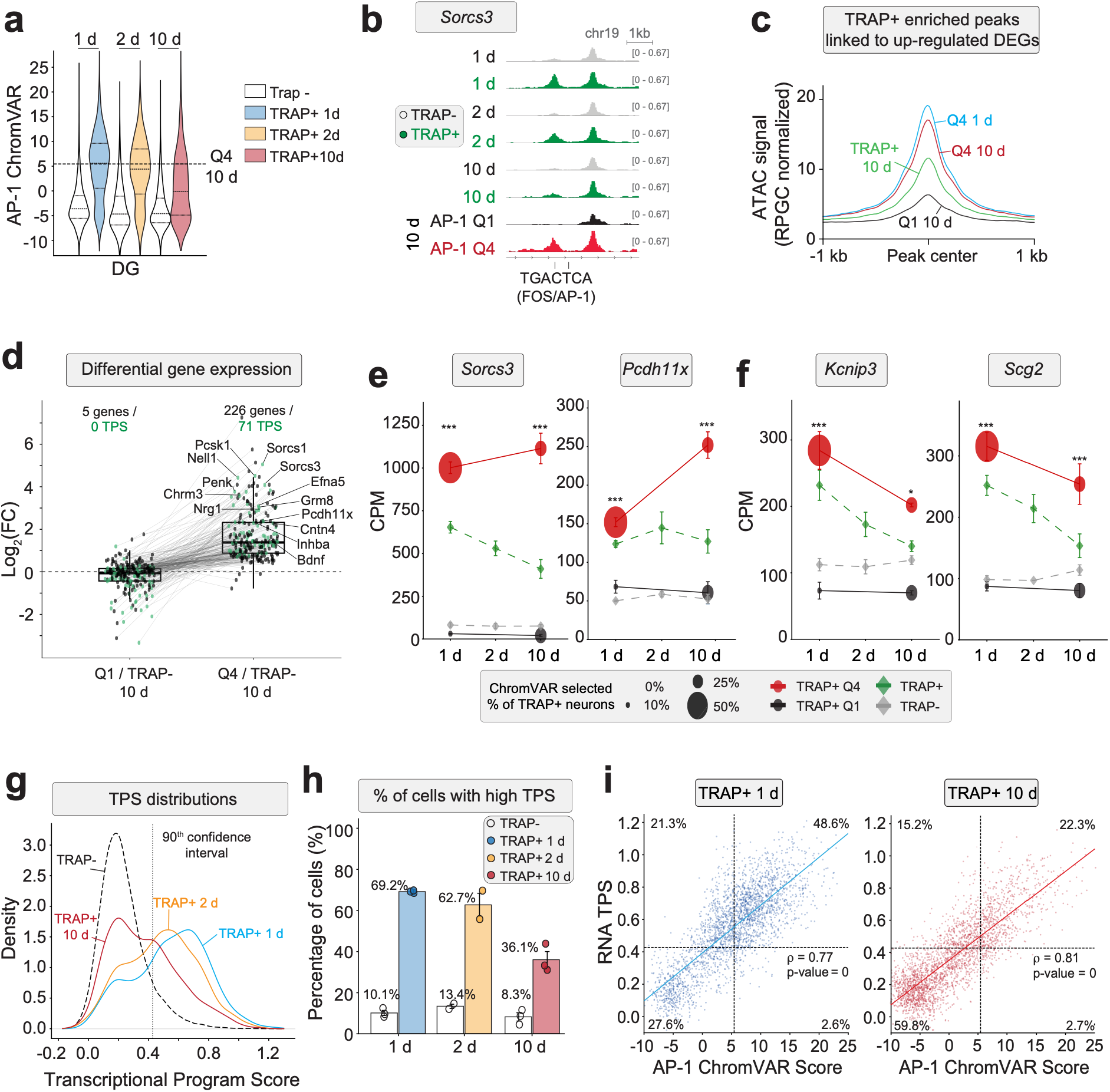
Heterogeneous maintenance of AP-1-associated chromatin and transcriptional states in DG ensemble neurons. a) Violin plot of AP-1 ChromVAR accessibility score distribution across TRAP+ and TRAP-DG neurons over time. Dashed line indicates cells contributing to Q4 at ten days. b) Example genome browser tracks of chromatin accessibility for activity-dependent and constitutive regulatory elements of TRAP-(grey) and TRAP+ (green) DG neurons over time. Selection for neurons in either AP-1 Low (Q1 black) or AP-1 High (Q4 red) quartiles is displayed. c) Aggregate plot showing peak accessibility around the peak center for TRAP+-enriched peaks linked to up-regulated DEGs in the DG. Accessibility is displayed for either all TRAP+ neurons at ten days (green), or either AP-1 Q1 at ten days (black), AP-1 Q4 at one blue or ten red days. Cells of Q4 at one day are derived from the cutoff based on Q4 at ten days as depicted in a). d) Box plots of mean Log_2_ Fold Change (DEseq2 on pseudobulked Quartile-cells vs 100 bootstrapped iterations of matched numbers of randomly selected TRAP-cells, median adjusted p-value from Wald test with Benjamini-Hochberg correction and mean Log2 Fold Change, Upregulated DEGs = median Padj<0.05 and Log2FC>0) for differential gene expression between TRAP-neurons and AP-1 Low (Q1) or AP-1 High (Q4). Numbers of differentially expressed genes are indicated, genes belonging to the Transcriptional program score (TPS, explained in g)) are highlighted in green. Lines indicate the change of a given gene between the two comparisons. e) Counts per million (CPM) of two persistent example genes, *Sorcs3* and *Pcdh11x*, showing their expression in either all TRAP+ neurons (green), all TRAP-neurons (black), ChromVAR-High TRAP+ (red) or ChromVAR-Low TRAP+ (black) neurons. Asterix indicate significance when TRAP+ Q4 neurons are compared to a randomly selected matched number of TRAP-cells (DESEq2 Wald test with Benjamini-Hochberg correction, *** < 0.001, * < 0.05) f) Counts per million (CPM) of transient example genes, *Kcnip3* and *Scg2*, showing their expression in either all TRAP+ neurons (green), all TRAP-neurons (black), ChromVARHigh TRAP+ (red) or ChromVAR-Low TRAP+ (black) neurons. Asterix indicate significance when TRAP+ Q4 neurons are compared to a randomly selected matched number of TRAP-cells (DESEq2 Wald test with Benjamini-Hochberg correction, *** < 0.001, * < 0.05) g) Density plot depicting the distribution of Transcriptional Program Scores (TPS) per cell for TRAP-neurons (black), or TRAP+ neurons at one day (blue), two days (orange) and ten days (red) post cFC. TPS represents the per-cell cumulative enrichment of the gene module computed from the top 100 TRAP+ up-regulated DEGs at one day. Dotted line depicts threshold for classification as TPS High and requires cells to be above the 90^th^ percentile among all TRAP-DG neurons. h) Percentage of neurons exhibiting a High TPS from either TRAP– or TRAP+ neurons at each time point. Bar plot displays mean ± SEM, circles indicate individual multiome replicates. i) Correlation between TPS and AP-1 ChromVAR score of TRAP+ neurons at one (blue) or ten (red) days. Spearman’s rank correlation coefficient (r), and percentage of neurons in each quadrant are indicated.

To better understand the nature of heterogeneity within the memory ensemble at ten days post cFC, we binned TRAP+ DG neurons into quartiles (Q1 = AP-1 low, Q4 = AP-1 high) based on their AP-1 ChromVAR scores and re-evaluated the accessibility of the previously defined DEGlinked DARs. DEG-linked DARs in Q4 neurons at ten days exhibited accessibility similar to that of Q4 neurons at the one-day timepoint, and were more accessible than the overall ten-day TRAP+ population or Q1 neurons at ten days (**Fig. 4b-c**). This enhanced maintenance of accessibility in Q4 neurons was most evident at DARs that we previously classified as persistent, and could also be detected at previously defined transient peaks, albeit to a lower extent (**Fig. S6a**). Thus, differences in AP-1 ChromVAR scores, potentially driven by differences in the level of long-lived AP-1 family members such as FOSB, reflect markedly distinct chromatin states at regulatory elements linked to the ensemble-specific gene expression program. This finding suggested that there may be significant heterogeneity in the gene expression programs expressed by ensemble neurons ten days following cFC and raises the possibility that the neurons with high AP-1 ChromVAR scores might be the subset of neurons that mediate memory retrieval.

### AP-1 motif accessibility predicts long-term ensemble transcriptional states

To determine if heterogeneity in AP-1-associated chromatin accessibility leads to distinct transcriptional states within DG ensemble neurons, we compared the gene expression programs of TRAP-neurons with TRAP+ Q1 or TRAP+ Q4 cells. This analysis revealed that the Q4 subset of TRAP+ neurons at ten days remains transcriptionally distinct from TRAP-neurons, both in the number of DEGs and in the magnitude of their expression (**Fig. 4d**, 226 significant genes, **S6b**). By contrast, the gene expression programs of Q1 TRAP+ neurons were largely indistinguishable from those of TRAP-cells (**Fig. 4d**, 5 significant genes). Notably, genes classified as persistent at the population level, including *Sorcs3* or *Pcdh11x*, remained stable or even increased their level of expression within the Q4 subset of neurons, despite the modest decline in expression detected when TRAP+ neurons were analyzed in aggregate (**Fig. 4e**). Transiently expressed genes, such as *Kcnip3* and *Scg2*, were also found to maintain their differential gene expression pattern at ten days when the analysis was restricted to Q4, although this effect was less pronounced than that seen for persistently expressed genes (**Fig 4f**). These findings suggest that the attenuation of ensemble-specific gene expression observed in memory ensemble neurons over time is due to heterogeneity within the TRAP+ population, and that sustained AP-1 DARs within a subset of TRAP+ neurons drive the persistent expression of plasticity genes within this subset of the memory ensemble.

To further examine the link between AP-1-dependent chromatin state and the persistence of gene expression over time, we calculated per-cell transcriptional program scores (TPS) representing a cell-level estimate of the relative maintenance of the ensemble gene expression state. This TPS calculation was derived for each TRAP+ and TRAP-DG neuron based on the level of expression of the 100 most differentially-expressed memory ensemble genes at one day post-cFC using gene modules computed with Scanpy^43^. The majority (56 genes) of the 100 most differentially expressed ensemble genes at one day also display persistent expression at ten days post-cFC. Mirroring our chromatin findings, we observed significant heterogeneity within the TRAP+ population using this TPS metric, with ∼69% of TRAP+ DG neurons displaying high TPS one day following cFC, whereas the remaining ∼31% exhibited a low TPS that was similar to that of TRAP-cells (**Fig. 4g, h**). Although the fraction of TRAP+ neurons with a high TPS declined over time, a substantial subset (∼36%) retained a high TPS at ten days post cFC (**Fig. 4h**). Thus, while ensemble gene expression signatures trend towards those of TRAP-DG neurons over time when analyzed in aggregate, a distinct identifiable subset of TRAP+ neurons preferentially retains persistent DEG expression as far out as ten days post cFC. Consistent with this, 71 out of the 100 genes in our TPS significantly increased their expression when ten-day Q4 was compared to TRAP-neurons, whereas no TPS genes in Q1 reached significance (**Fig. 4d, S6b** comparison Q4 to Q1 at ten days, TPS genes labeled in green). Notably, ChromVAR and the TPS were strongly correlated on a per-cell basis, with Spearman correlation coefficients of 0.77 and 0.81 at one and ten days, respectively (**Fig. 4i, S6c**). Taken together, these findings suggest that sustained accessibility of AP-1-associated regulatory elements promotes transcriptional persistence in ensemble neurons. The apparent degradation of the ensemble gene expression program at the population level may therefore reflect a progressive drop-out of a subset of cells from the memory ensemble over time, as their AP-1 chromatin state resets to that of TRAP-DG neurons.

### *Sorcs3* expression as a predictor of DG neuronal memory ensemble reactivation

A hallmark of neurons within memory ensembles is their preferential reactivation of the *Fos* mRNA during recall. However, the fraction of neurons within the ensembles that re-express Fos is typically quite low (∼10-30%)^1,16^. We hypothesized that this small group of TRAP+ neurons might be a molecularly distinct subset of TRAP+ neurons characterized by a high TPS. To begin to address this hypothesis, we used expression of *Sorcs3* mRNA as an indicator of TRAP+ neurons with a high TPS. We then trapped ensemble neurons during cFC as described above and returned the mice to their home cage for either one or ten days. Animals were then either re-exposed to the training context without shock to induce memory retrieval (“5′ recall”) or maintained in the home cage (“No recall”) (**Fig. 5a**). Importantly, to avoid potential downstream gene induction following acute FOS re-expression, we harvested hippocampal tissue 25 minutes after recall. We then performed FISH for *Fos* mRNA to identify reactivated neurons, and *Sorcs3* mRNA, to identify the subset of TRAP+ neurons with a high TPS.

**Figure 5:**
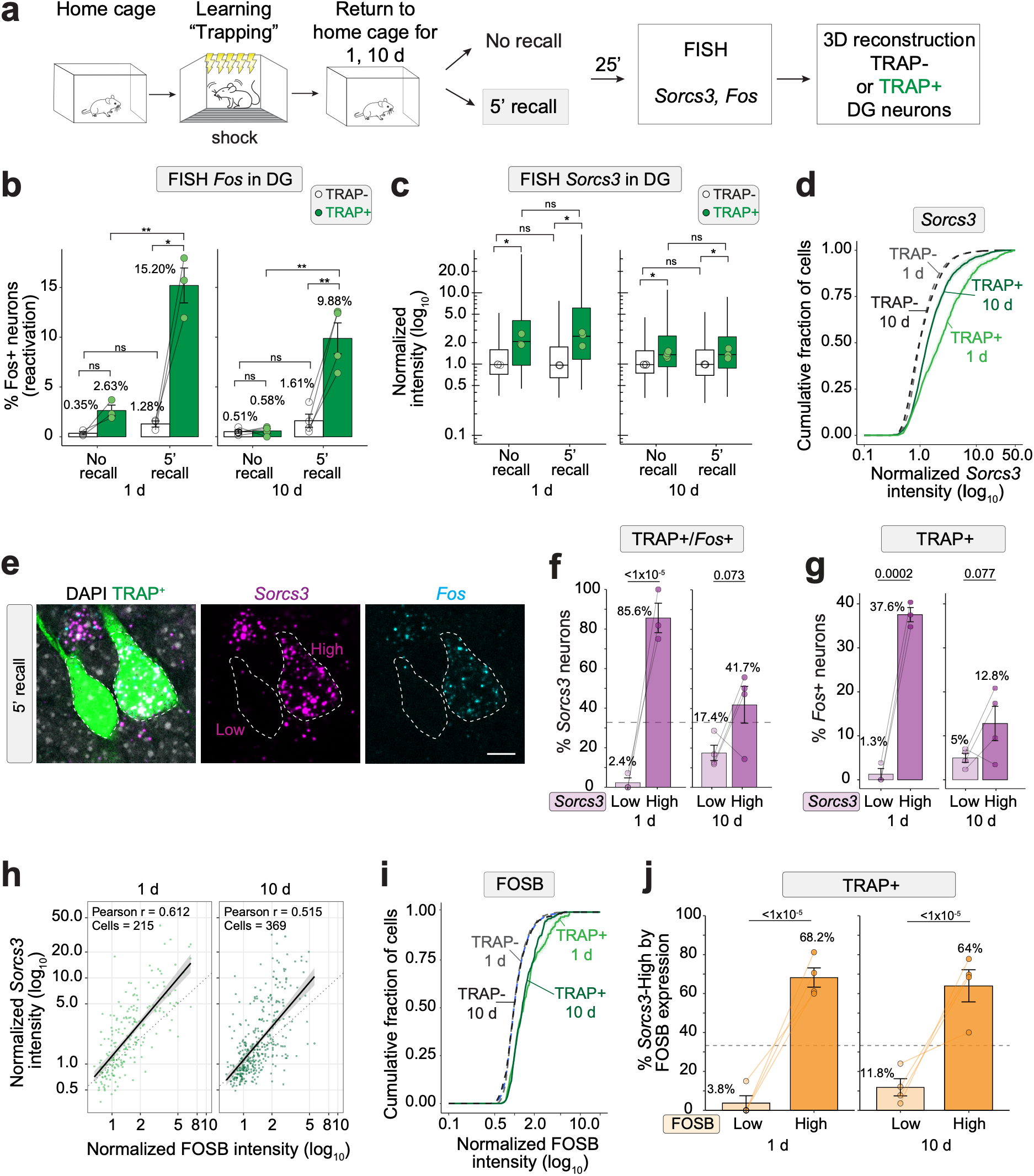
*Sorcs3* mRNA expression indicates preferential reactivation of dentate gyrus memory ensemble neurons. a) Experimental design for assessing ensemble neuron reactivation after five-minute memory recall at one– or ten-days post-cFC. b) Percentage of FOS+ neurons with or without five-minute memory recall at one– or ten-days post-cFC. Small circles indicate independent biological replicates. Mean ± SEM is indicated. One day no recall: N = 3 animals, TRAP+: 502 cells / TRAP-: 11444 cells; One day recall: N = 3 animals, n = TRAP+: 293 cells / TRAP-: 6180 cells; Ten days no recall: N = 4 animals, n = TRAP+: 757 cells / TRAP-: 10486 cells; Ten days recall: N = 4 animals, TRAP+: 577 cells / TRAP-: 10529 cells. Significance assessed using ANOVA with posthoc t-testing as described above. TRAP⁺ vs. TRAP⁻ differences were significant only in the recall condition at both timepoints (1 day, Padj = 0.017; 10 days, Padj = 0.006). Within TRAP⁺ cells, NR vs. REC comparisons also significant at both timepoints (one day Padj = 0.002; ten day Padj = 0.001). Normalized Log_10_ intensity values of FISH puncta per cell for *Sorcs3* mRNA with or without five-minute recall at one– or ten-days post-cFC. Box plots depict median, One day no recall: N = 3 animals, n = TRAP+: 502 cells / TRAP-: 11444 cells; One day recall: N = 3 animals, n = TRAP+: 293 cells / TRAP-: 6180 cells. Ten days no recall: N = 4 animals, n = TRAP+: 757 cells / TRAP-: 10486 cells; Ten days recall: N = 4 animals, n = TRAP+: 577 cells / TRAP-: 10529 cells. Note that data for ‘no recall’ have also been displayed in Fig. 1j. ANOVA with post hoc t-tests: TRAP⁺ vs TRAP⁻ cell comparisons (one day NR Padj = 0.049, one day REC Padj = 0.041, 10 d NR Padj = 0.011, ten day REC Padj = 0.03). Within TRAP⁺ comparisons of NR to REC NS at both timepoints (one day TRAP+ Padj = 0.575, ten day TRAP+ Padj = 0.95) c) Cumulative distribution of the Log_10_ normalized Sorcs3 FISH intensity values for cells after memory recall. Dashed lines TRAP-neurons: one day (grey), ten days (black) post cFC. Solid lines TRAP+ neurons: one day (light green), ten days (dark green) post-cFC. d) Representative FISH image of DG neurons after five-minute memory recall one day post cFC. DAPI (grey), TRAP+ neurons (green), *Sorcs3* mRNA (magenta), *Fos* mRNA (cyan). Scale bar = 10 µm. e) Percentage of *Sorcs3* neurons in TRAP+/Fos+ reactivated DG neurons. *Sorcs3* Low or High tertiles are displayed. One day recall: N = 3 animals, n TRAP+: 47 cells, Ten day recall: N = 4 animals, n TRAP+: 57 cells. Horizontal dotted line indicates expected percentage of neurons reactivated by chance. P-values derived through parametric assessment of likelihood of observed distribution across tertiles relative to null expectation of equal division using Monte Carlo goodness-of-fit test (one day p-value < 1×10^-5^, ten day p-value 0.073) f) Percentage of *Fos*+ neurons in TRAP+ DG cells, grouped based on their *Sorcs3* expression (Low or High). One day recall: N = 3 animals, TRAP+: 293 cells; Ten day recall: N = 4 animals, TRAP+: 577 cells. Based on an a priori hypothesis that higher Sorcs3 expression is a marker of the overall chromatin/transcriptional state that supports neuronal reactivation, one-sided paired t-tests were used to assess whether Fos positivity was greater in Sorcs3-high compared to Sorcs3-low TRAP+ cells (1 day: p = 0.0002; 10 days: p = 0.077). g) Correlation plot between FOSB protein expression and *Sorcs3* mRNA levels in DG neurons at one– or ten-days post-cFC in the absence of memory recall. Pearson r correlation coefficient and numbers of cells analyzed are indicated. h) Cumulative distribution of the Log_10_ normalized FOSB protein intensity values for cells without memory recall. Dashed lines TRAP-neurons: one day (grey), ten days (black) post-cFC. Solid lines TRAP+ neurons: one day (light green), ten days (dark green) postcFC. i) Percentage of *Sorcs3* mRNA-High cells grouped based on their FOSB protein expression (Low or High). One day no recall: N = 4 animals, n = TRAP+ 215 cells; Ten day no recall: N = 4 animals, n = TRAP+ 369 cells. Horizontal dotted line represents percentage of neurons detected by chance. Monte Carlo goodness-of-fit test against null expectation of equal distribution (one day p-value < 1×10^-5^, ten day p-value < 1×10^-5^)

As expected, we detected very few cells that express the *Fos* mRNA in mice that were not exposed to the recall stimulus (2.6% at one day and 0.6% at ten days) and observed re-activation rates of ∼15% and ∼10% of TRAP+ neurons at one– and ten-days post cFC, respectively (**Fig. 5b**), similar to previously reported reactivation rates^1,16^. Reflecting the molecular heterogeneity of the TRAP+ memory ensemble that we observe at all three remote timepoints (**Fig. 4a, g**), the level of *Sorcs3* mRNA was variable across TRAP+ neurons (**Fig. 5c, d**). To determine whether a high level of *Sorcs3* mRNA expression is associated with DG ensemble reactivation upon memory recall, we classified TRAP+ cells according to their *Sorcs3* mRNA expression level (*Sorcs3*-Low = Bottom Tertile, *Sorcs3-Mid* = Middle Tertile, *Sorcs3*-High = Top-Tertile, **Fig. S7a**) and examined their contribution to the reactivated (TRAP+/*Fos+*) population. One day post cFC, 85.6% of the reactivated TRAP+ *Fos+* neurons were *Sorcs3*-High while only 2.4% were *Sorcs3*-Low. Ten days post cFC, 41.7% of TRAP+ *Fos+* neurons were *Sorcs3*-High while 17.4% were *Sorcs3*-Low (**Fig. 5f**), suggesting that memory recall preferentially engages neurons with high *Sorcs3* expression, especially during early phases of memory consolidation. We further asked whether *Sorcs3* levels are linked to the likelihood that TRAP+ neurons are reactivated. Expanding our analysis to the entire TRAP+ population revealed that *Sorcs3*-High TRAP+ neurons were on average 28-fold more likely to reactivate (express the *Fos* mRNA) than *Sorcs3*-low TRAP+ neurons one day postcFC (37.6% vs. 1.3% p = 0.0002). At the 10-day timepoint, the animal-level average reactivation rate remained higher in Sorcs3-high cells compared to Sorcs3-low cells (12.8% vs. 5%; ∼2.5-fold; **Fig. 5g, Fig. S7b**), although this trend did not reach statistical significance (p = 0.077). Altogether, these findings indicate that elevated *Sorcs3* mRNA expression (e.g. a high TPS) marks a subset of DG ensemble neurons that exhibit a markedly increased likelihood of reactivation upon memory recall.

Finally, we asked if a high level of *Sorcs3* mRNA expression, and thus the likelihood of neuronal reactivation during recall, is correlated with a high level of FOSB protein expression prior to recall. Such a correlation would provide support for the hypothesis that the sustained expression of the FOSB protein promotes the long-lasting expression of *Sorcs3* mRNA, and other plasticity associated mRNAs, and in this way mediates DG ensemble neuron reactivation and memory retrieval. Towards this end, we quantified the co-localization of FOSB protein and *Sorcs3* mRNA in DG neurons in the absence of recall. This analysis revealed a strong correlation between the level of FOSB protein and *Sorcs3* mRNA expression in TRAP+ neurons at one– and even tendays post-cFC (Pearson correlation coefficient r=0.612, r=0.515, respectively, **Fig. 5h**). The range in the intensity of signal in FOSB+ cells allowed us to bin FOSB+/TRAP+ cells into intensity tertiles in the same manner as we did for Sorcs3. Classifying cells according to their FOSB protein intensity and examining their respective rates of *Sorcs3*-High mRNA levels revealed that 68.2% and 64% of TRAP+/FOSB-High neurons were *Sorcs3*-High at one– or ten-days post-cFC, respectively. By contrast, less than 4% and 12% TRAP+/FOSB-low neurons were *Sorcs3*-High at one– and ten-days, respectively, after cFC (**Fig. 5i, j, Fig. S7c**). Taken together with our chromatin accessibility and motif analyses, these findings suggest that long-lasting FOSB expression sustains the expression of a battery of memory ensemble-associated genes, including *Sorcs3*, for weeks following cFC in a core subset of TRAP+ DG neurons that are then poised for reactivation upon memory retrieval.

## DISCUSSION

A central question in neuroscience concerns how transient stimuli are stably encoded into the long-lasting circuit changes that underlie learning and memory. While IEGs such as *Fos* have been suggested to regulate early phases of memory formation during the first day poststimulation, the mechanism by which these short-lived signals are translated into persistent, ensemble-specific plasticity that can support lifelong memories remains poorly understood. In this study, we uncovered multiple functional changes, spanning circuit-level rearrangements, chromatin remodeling and transcriptional dynamics, that likely contribute to how DG neuron memory ensembles are shaped and maintained in mice weeks after the stimulation event, allowing for the retrieval of memories at these distant times. Instead of the canonical model in which IEG induction triggers a wave of late-response gene expression that returns to baseline within hours^44^, we find that within memory ensembles, IEG expression triggers a long-lasting imprint on neuronal subtype-specific gene regulatory elements that leads to transcriptional responses that extend out to ten days and likely longer. These sustained gene expression programs encode numerous modulators of synaptic and intrinsic plasticity, raising the possibility that these signatures are not merely correlates of ensemble identity but actively reshape ensemble function over time. The identification of the genes that are persistently expressed in memory ensembles provides a roadmap for further mechanistic understanding of the process by which neuronal ensembles encode long-lasting memories.

In line with this model, persistent DEGs in DG ensemble neurons include multiple regulators of excitatory synaptic transmission, such as *Sorcs1, Sorcs3* and *Grip1*, that have been linked to the regulation of AMPAR signaling^26,27,45^. This suggests that sustained activation of these genes may contribute to the persistent increase in mEPSC frequency in ensemble neurons observed at two and ten days post-cFC. By contrast, genes that are more readily detected at one day in DG neurons following cFC include the neuropeptide-encoding genes *Scg2* and *Bdnf,* and the GABAR subunit-encoding gene *Gabrd* in the DG, each of which regulate inhibitory synaptic transmission in the hippocampus and cortex^29,30,46–48^. Thus, genes such as *Scg2* may contribute to this dynamic change in inhibition. The decrease in intrinsic excitability at two days post-stimulation coincides with, and may be due to, transient up-regulation of *Kcnip3*, a voltage-gated potassium channel auxiliary component that is known to dramatically reduce neuronal excitability in an activitydependent manner by altering potassium currents^31^. Intriguingly, we also detect a multitude of genes that contribute to axonal guidance or synapse specification such as *Nrxn3* or *Pcdh11x*^49–51^. The dynamic nature of gene expression of these cell adhesion molecules might further support the stability of synaptic connectivity onto DG ensemble neurons, as well as their outputs to downstream areas such as CA3 that might be required for the transfer of the memory within and outside the hippocampus. Taken together, these findings suggest that activity-dependent transcription coordinates a shift in both the synaptic and intrinsic excitability properties of ensemble neurons, leading to memory encoding and ultimately memory retrieval. In the context of the neuronal circuit, this activity-dependent shift in intrinsic excitability and synaptic balance likely reconfigures how ensemble neurons gate and route incoming information, enabling the DG to transition from initial memory encoding to a more stable, pattern-separated representation of memory over time.

Understanding how memory ensembles evolve to support neuronal circuit adaptations during memory consolidation requires resolving the gene regulatory and transcriptional mechanisms selectively engaged in neurons actively participating in the memory ensemble (1-2% of hippocampal neurons). Reliable identification of these mechanisms therefore required paired chromatin accessibility and transcriptomic profiling with single-cell resolution from a sufficiently large number of neurons. By purifying a substantially greater number of memory ensemble neurons than has previously been achieved, profiling these neurons at various times post-cFC with single-cell multiomic resolution, and controlling for identical physiological experiences by comparing TRAP+ to TRAP-neurons from the same animals, we were able to obtain for the first time a sufficiently rich single nucleus dataset to identify the mechanisms that underly the dynamic chromatin states within the neuronal ensemble. Our work revealed plasticity-relevant transcriptional programs that likely control physiological changes within the ensemble that are essential for memory encoding. Furthermore, this depth was essential to resolve the molecular heterogeneity within individual cell types of ensemble neurons, which represents a previously unrecognized layer of molecular regulation shaping the evolution of memory ensembles over time.

The dynamic features and heterogeneity of memory ensembles that we have uncovered reinforce the view that ensemble formation is not an instantaneous event, but a prolonged process of maintenance and maturation that unfolds over hours, days, and weeks as memories undergo systems consolidation^1,2^. The changes in transcriptional states over time that we document here for DG granule cells may constitute a fundamental mechanism by which memory ensembles undergo temporal refinement, allowing these ensembles to remain both stable and adaptable. Cells that revert their gene expression profiles may become available to encode new experiences, whereas cells that maintain a new transcriptional and electrophysiological state may continue to participate in the ensemble, thereby balancing flexibility with persistence in memory representation.

The analysis of gene regulatory elements within memory ensembles has provided additional mechanistic insight into the evolution and heterogeneity of TRAP+ DG neurons over time. FOS and its related family members are not merely markers of the memory ensemble. Rather, the presence of the AP-1 site within 80% of open regions in TRAP+ neurons at ten days strongly suggests that FOS and/or its related family members are key mediators of memory consolidation. Somewhat surprisingly, our chromatin accessibility data indicate that the AP-1 motif and the factors it binds are not just required for the transient activation of gene expression in memory ensemble neurons but instead function as a central determinant of sustained chromatin opening that promotes long-term transcriptional stability in a subset of cells. We propose that these AP-1 motif-containing regulatory elements are initially bound by transiently expressed FOS, which is known to recruit members of the BAF chromatin-remodeling complex to trigger the opening of chromatin and the activation of gene transcription. In this scenario, FOSB may function in a manner similar to FOS and could contribute to the initial opening of chromatin and the activation of gene transcription. Previous work has demonstrated that repeated behavioral training or cocaine stimulation leads to a gradual increase in the expression of a long-lived short form of FOSB (ΔFOSB) with a half-life of eight days that has been suggested to contribute to learning^37–39,52–54^. Our observations extend these findings and suggest a model in which FOS and the longform of FOSB decay away within hours after a very brief stimulus (cFC), while long-lasting ΔFOSB protein expression keeps crucial regulatory elements in an active state to prolong gene expression in ensemble neurons, supporting long-term memory. Both FOS and the long form of FOSB are known to activate transcription as heterodimers with members of the JUN family, and thus it is likely that one or several activity-dependent JUN family members also are involved in the initial activation of gene expression during the formation of memory ensembles. Moreover, ΔFOSB is known to form homodimers, and heterodimers with JUND, either of which may be responsible for maintaining activity-dependent gene transcription in memory ensembles for many weeks following cFC.

A number of prior studies have highlighted a role for the promoter-bound transcription factor CREB in driving the expression of *Fos* and other IEGs, as well as a link between CREB levels in the lead up to learning and neuronal excitability regulation – positioning this factor as a critical determinant of ensemble allocation^4^. However, while a large fraction of the DEGs we have identified possess CREB motifs within their promoters **(Fig 3d, S4c)**, this motif is equally common at accessibility-matched non-DEG promoters. Thus, it is unlikely that CREB alone would be sufficient to specify the persistent transcriptional states of ensemble cells. It strikes us that the initial FOS or full-length FOSB binding at ensemble-specific AP-1 motif-containing enhancers could amplify CREB’s action at promoters of ensemble-specific DEGs, while long-lived ΔFOSB might maintain these targets in ensemble neurons over time.

We speculate that this mechanism may be used widely in many regions of the brain that encode various long-term memories and behaviors in response to brief or repeated stimuli^55,56^. Brain regions such as the medial prefrontal cortex, amygdala or thalamus form ensembles with markedly different temporal properties from those in the hippocampus^1,10,16,22,56–59^. However, it is possible that the greater stability of ensemble activity reported in these areas nevertheless reflects a similar mechanism of persistence and heterogeneity in ensemble-wide gene-expression programs and neuronal circuit refinement that we describe for the hippocampus. Furthermore, while the reduction in the responsiveness of training-tagged ensemble-neurons is thought to be a key element of ensemble evolution over time, there is evidence that the integration of additional neurons to the ensemble throughout later periods of consolidation or during recall may contribute to memory retrieval as well. Our approach was tailored to capture the evolution of the population of neurons active during cFC training and was able to identify neurons that drop out of the ensemble over time. However, it is likely that incorporation of new neurons into the ensemble also occurs, and it will be important for future studies to compare and contrast the mechanisms at play in these alternative trajectories of ensemble cells.

Our data suggests that ensemble neurons can exhibit distinct functional, transcriptional, and regulatory states at the time of reactivation due to cellular heterogeneity. Thus, their state at the time of reactivation may selectively tailor transcriptional remodeling and plasticity that could be critical for further neuronal ensemble refinement and evolution in various behavioral paradigms. Moreover, our discovery of the persistent transcriptional program, exemplified by the high rate of *Sorcs3* expression in a subset of reactivated DG ensemble neurons, provides a molecular handle to identify and functionally characterize the subset of ensemble neurons critical for the recall of memories. In line with this, the characterization of activity-dependent gene regulatory elements of genes such as *Sorcs3* may further allow for the generation of novel tools to selectively label and manipulate these critical subsets of neurons that express a memory trace over days to weeks after the initial experience. In addition, our identification of activity-dependent and cell classspecific gene expression programs provides a rich resource for the memory field and will enable researchers to dissect in diverse neuronal subtypes the precise functions of these programs for memory consolidation. Together, our findings reveal a previously underappreciated layer of ensemble complexity at the epigenetic and transcriptional level, offering a mechanism for how neurons selectively engage with, and mature within, the ensemble over time.

## METHODS

### Mice

Mice were handled and subsequent experiments were performed according to protocols approved by the Harvard University Standing Committee on Animal Care and were in accordance with federal guidelines. Wild-type C57/BL6 (JAX 000664), FosTrap2-CreERT2 mice (JAX 030323), or floxed Fos, FosB, JunB (FFJ) triple recombinant mice^60^ were housed in a 12 hr light/dark cycle with ad libitum access to food and water. Temperatures range from 20-23°C, with humidity between 35-65%. All experiments were performed during the light cycle. For all sequencing studies, male mice were used to avoid the confounding effects of estrogen cycles observed in females, but for further histology, molecular biology and electrophysiological experiments both males and female mice were used. In these follow-up experiments no differences between sexes were observed.

### Stereotaxic injection

Adult mice (8-12 weeks of age) used for behavioral, genomic and molecular characterization were anesthetized using isoflurane inhalation on a stereotaxic injection frame (Kopf Model 1900 Stereotaxic Alignment System) and body temperature was maintained at 37°C using a heat pad. After ensuring that mice were deeply anesthetized, fur around the scalp area was removed using a shaver and sterilized with three alternating wipes with betadine and 70% ethanol. To target the whole hippocampus as best as possible two burr holes per hemisphere were drilled 1) AnteriorPosterior, AP: –2.2mm, Medial-Lateral, ML ± 2mm, Dorsal-Ventral, DV –1.6mm and 2) AnteriorPosterior, AP: –2.75mm, Medial-Lateral, ML ± 3mm, Dorsal-Ventral, DV –2.8mm. For electrophysiological experiments, stereotaxic injection was performed either at postnatal day P2834 for 2 days or P21-26 for 9/10 days after fear learning, allowing for characterization at the same developmental age between P40 – 46. Coordinates for young adults were AP –2.4mm, ML ±1.8 and DV –1.6mm. Adeno-associated viruses (AAVs) were slowly injected using a glass pipette pulled on a Sutter P-2000, and pipette was left in place for 5 min to allow for viral spreading before slow removal. All animals were given postoperative analgesic by subcutaneous injection of Buprenorphine SR/ER (0.5-1mg/kg) or topical application of Zorbium transdermal buprenorphine at 40 mg/kg and monitored for four consecutive days after the surgery for signs of pain or infection.

### Plasmids, Viral vectors and titers

For labeling of nuclei for fluorescence-activated nuclear sorting (FANS) AAV2/9-flex-KASH-EGFP (Addgene #154373, AAV produced by Canadian Neurophotonics Platform Viral Core) was injected at a titer of 3.7 x 10^11^ gc/mL per injection site into FosTrap2-CreERT2 mice. Viral injection for whole-cell labeling and subsequent confocal imaging was performed similarly using AAV2/1hSyn-dio-mCherry (Addgene #50459). For electrophysiological characterization of TRAP+ dentate gyrus granule cells, we generated an AAV construct containing a floxed nuclear blue fluorescent protein (mTagBFP2) driven by the human synapsin promoter (AAV-dio-nBFP) which was packaged into AAV2/9 by the Boston Children’s Hospital Viral Core injected at 1×10^12 gc/mL. The AAV expressing Cre recombinase fused with GFP and was injected at a titer of 1.1×10^11^ gc/mL.

### Fear conditioning paradigm

Five days after stereotaxic surgery mice were single housed and brought to the temperature and humidity-controlled behavior room. Mice were subsequently handled by the experimenter multiple times once daily for three days to habituate them to handling. To label neurons with heightened neuronal activity, mice mildly anesthetized with isoflurane were injected with tamoxifen (4OHT, Sigma Aldrich #H6278) at 50mg/kg intraperitoneally 30 min prior to the fear conditioning training session. Mice were placed in a Habitest Modular System FreezeFrame Fear conditioning setup (Coulbourn) containing a stainless-steel shock grid. Mice were allowed to freely explore the novel context for three minutes, after which five shocks were delivered every 30 seconds with a 1 second electric pulse between 0.2-0.8 mA. Mice were kept in the conditioning chamber for one additional minute after delivery of the last shock to assess freezing behavior of the animal as an indicator of appropriate training. For memory recall, mice were placed back into the training arena without delivery of foot shocks for five minutes either one or ten days after training. The behavior of the animal was monitored and recorded using a video-based tracking system throughout training and recall sessions. After each session mice were returned to their home cage. In between animals, the conditioning chamber and grid were cleaned with 70% ethanol.

### Preparation of 4OHT

4-Hydroxytamoxifen (Sigma-Aldrich, H6278) was dissolved in 450 µL ethanol with vortexing to facilitate solubilization and heated at 37c for 5 minutes. Sunflower oil and castor oil were mixed at a 4:1 ratio, and the drug solution was combined with an equal volume of this oil mixture and vortexed. The mixture was heated at 37c for an additional 5 minutes and then transferred to a vacuum centrifuge for 30 minutes to evaporate the ethanol, leaving the drug suspended in the oil. Additional oil mixture was added to achieve a final concentration of 6.67 mg/mL. Mice were administered the final suspension by IP injection at a dose of 50 mg/kg.

### Single-nuclei Multiome nuclei preparation

To account for batch effects during library preparations or sequencing, one biological replicate of all three timepoints (1, 2 and 10 days) were collected within three days. Biological replicates for all timepoints were collected over the course of multiple months, further highlighting reproducibility. Mice were deeply anesthetized with isoflurane, decapitated, and brains were placed on an ice-cold metal block. Hippocampi were extracted under a dissection microscope based on anatomical features and immediately processed for nuclei preparation: Each hippocampal hemisphere of each animal was processed separately. Tissue was homogenized in 2 mL Homogenization buffer in a 7 mL glass dounce with 10 strokes loose and 10 strokes tight, after which 64 µL of 5% Igepal was added before douncing 8x with a tight pestle. The homogenate was filtered through a 40 µm strainer, transferred to a 2 mL DNA low-bind tube, and centrifuged at 500g for 10 min at 4°C in a swing bucket. Supernatant was discarded and the pellet was resuspended in 500 µL of Homogenization buffer containing Anti-NeuN-Alexa Fluor-555 Conjugate (Sigma Aldrich #MAB377A5) at a 1:500 dilution to sort for NeuN-positive nuclei and incubated on ice for 15 min. Subsequently, nuclei were transferred into a 1.5 mL Protein-lowbinding tube and centrifuged in a swing bucket at 500g at 4°C. During this time, 1.5 mL Proteinlow-binding tubes were coated with 300 µL of 30% BSA (RNAse free, Sigma A-8577) for 5 min at room temperature. The pellet was resuspended in 1.2 mL of Homogenization buffer and nuclei were stained with 7AAD (Sigma, SML1633, 1:120 dilution).

Nuclei were fluorescence-activated nuclei sorted (FANS) using a Sony SH800Z cell sorter. Sorting was performed using a 70 µm microfluidic sorting chip at a sample pressure of 6 and event rates between 500–7,000 events per second. Nuclei were collected into 300 µL of Nuclei Collection Buffer. Events were first gated based on forward and side scatter (FSC-A vs SSC-A) to exclude debris, followed by doublet discrimination using FSC-A versus FSC-H to retain singlet nuclei. Nuclei were then identified based on 7-AAD DNA staining signal, and neuronal nuclei were enriched based on NeuN-555 signal. KASH-EGFP-positive nuclei were subsequently gated based on fluorescence intensity. EGFP gating thresholds were defined by comparing GFP signal from IP vehicle-injected x Flex-KASH-EGFP stereotax injected control animals in comparison to experimental 4OHT injected animals to establish baseline autofluorescence and background signal, and consistent gating boundaries were applied across experiments to ensure reproducibility.

For each biological replicate, micro-dissected hippocampi from 5-6 mice were pooled to obtain a total of 60-70.000 GFP-positive and GFP-negative nuclei. Importantly, nuclei were kept separately for each mouse and hemisphere and checked for GFP-positivity rate prior to deciding whether to include the animal in the final dataset. GFP-positivity rates for included animals ranged between 0.8 and 3.5% consistent with what has been reported in the literature.

After collecting nuclei into one tube, volumes were measured, and Digitonin and Tween-20 were added to a final concentration of 0.01% and 0.1%, respectively. Nuclei were incubated at room temperature for 10 min. Samples were centrifuged in a swing bucket at 500g for 10 min at 4°C and supernatant was removed, leaving ∼20-30 µL behind. Dilute Nuclei Buffer (250 µL) was added carefully along the side of the tube wall without disturbing the nuclear pellet, and samples were centrifuged at 500g for 10 min at 4°C in a swing bucket. Supernatant was carefully removed without disturbing the pellet, leaving ∼7 µL, and the pellet was resuspended in the remaining buffer. Nuclei were counted on a hemocytometer, and 16,000 nuclei were used as input for the tagmentation reaction and 10X Chromium controller loading following standard protocols from the 10X platform. After tagmentation and reverse transcription, samples were frozen at –80°C until samples from all three time points were acquired and library preparations could be performed together. The DNA and RNA libraries were generated according to the manufacturer’s recommended protocol (https://www.10xgenomics.com/support/single-cell-multiome-atacplusgene-expression). 10x multiome ATAC–seq and RNA-sequencing (RNA-seq) libraries were sequenced together on the Illumina NovaSeq X systems using paired-end 2×150 bp reads to a depth of around 30,000 reads per cell for each modality.

### Homogenization Buffer (HB)

20 mM Tricine-KOH pH=7.9, 250 mM Sucrose, 25 mM KCl, 5 mM MgCl_2_, add fresh on day of: 1 mM DTT (P2325 Life Technologies), 0.15 mM Spermine (S1141 Sigma), 0.5 mM Spermidine (S02116 Sigma), RNAse protector inhibitor 40 U/µL (#3335402001 Roche) and cOmplete Mini EDTA-free protease tablet (#1183617001 Roche)

### ATAC RSB Buffer

10 mM Tris-Hcl pH=7.4, 10 mM NaCl, 3 mM MgCl_2_

### Nuclei collection Buffer

for 1 mL use 904 µL of ATAC RSB Buffer, 70 µL 30% BSA (Sigma, A-8577), 1 µL 1 M DTT and 25 µL RNAse protector inhibitor

### Dilute Nuclei Buffer

50 µL Nuclei Buffer (20X from 10X Genomics kit), 1 µL 1 M DTT, 25 µL Protector RNAse inhibitor and 924 µL Nuclease-free H_2_O.

### snRNA-seq Analysis

Raw sequencing data were processed using Cell Ranger ARC (v2.0.2, 10x Genomics) under default settings. Reads were mapped to mouse mm10 genome, with the GENCODE vm23 GTF file as used in the BICCN atlas. We performed unsupervised clustering with RNA UMI counts based on the SCANPY analysis pipeline^43^ and best practice steps. In brief, we first filtered for lowquality nuclei by requiring ≥1,000 ATAC fragments and ≥200 genes detected per nuclei and < 5% mitochondria RNA reads. We then performed a first-pass clustering for quality control. The top 3,000 highly variable genes were identified for principal component analysis (PCA). Top 30 PCs were used to perform K-nearest neighbor (k=25) graph-based Leiden clustering (resolution=1). Putative multiplets were predicted using Scrublet,6 and Leiden clusters with mean doublets score > 0.2 were marked as doublets. While a large majority (87-92%) of cells generated from Cell Ranger ARC passed these filtering steps we found that two samples (the replicate 1 two-day Trap+ and Trap-samples) exhibited significantly worse RNA quality than all other samples and thus were both excluded from further analysis of their RNA modality, rendering the statistical power for gene expression analyses more limited at the two day timepoint compared to the one day and ten day comparisons. For the remaining 16 samples we then integrated our high-quality cells with hippocampal formation (HPF) cells selected from the whole-mouse brain atlas dataset from BICCN using Seurat (v4.0.0).2,7 Each dataset was normalized using SCTransform with regression of mitochondrial read percentage. The top 3,000 variable features were used to identify integration anchors with FindIntegrationAnchors (dims = 1:30), followed by data integration with IntegrateData. PCA was performed on the integration matrix, and the first 30 PCs were used to construct a shared nearest neighbor (SNN) graph and identify clusters via modularity optimization (FindNeighbors and FindClusters, resolution=0.4). We annotated 14 major cell types, and 33 subclass level cell types based on the integration results as the cell type nomenclature used in this study. To visualize clusters and annotations, we used scVI package8 (v1.2.0 n_hidden = 512, n_latent = 40, n_layers = 2, use sample as batch_key) to learn a latent space and perform UMAP embedding9 to generate the visualization coordinates for scatter plots.

#### DG Subclustering

Subclustering of dentate gyrus ensemble neurons was performed using Scanpy. Highly variable genes (n = 3,000) were identified from raw counts using the Seurat v3 method, and principal component analysis (PCA) was performed on the top 30 components. A k-nearest neighbor graph (k = 30) was constructed using these components, followed by Leiden clustering (resolution = 0.5), resulting in 9 DG subclusters. These clusters formed a largely continuous transcriptional landscape with enrichment of TRAP+ cells in specific regions of the embedding. Inspection of cluster-specific gene expression revealed that the smallest cluster (n = 440 cells; <1% of DG cells) exhibited high expression of canonical CA3 markers and, when analyzed jointly with CA3 populations, clustered more closely with CA3 neurons than with DG neurons. This cluster likely represented misannotated or contaminating CA3 cells that were retained during initial subclass annotation and was excluded from all downstream analyses.

### Pseudobulk differential expression analysis

We focused our differential expression analyses on five major neuronal cell types robustly represented in TRAP+ and TRAP-populations across all replicates: DG granule cells, cornuammonis (CA) 3 and CA1 excitatory neurons, as well as somatostatin (SST)– and parvalbumin (PV)-positive interneurons. For each biological sample, raw counts were summed across cells belonging to the same neuronal cell type, generating one pseudobulk count matrix per cell type, per biological sample. Differential expression between the resulting TRAP+ and TRAP− pseudobulk counts matrices was assessed independently within each timepoint using DESeq2, with replicate included as a covariate. As TRAP+ and TRAP− populations were derived from the same pooled hippocampal tissue within each replicate, this design controlled for experimentspecific variability while testing for differences attributable to TRAP status. Normalized expression values were obtained using DESeq2 size-factor normalization. For visualization, including dot plots, pseudobulk count matrices from selected cell types were additionally merged and jointly normalized to generate globally comparable expression values across samples and cell types. Statistical significance was assessed using Wald tests implemented in DESeq2, and genes were considered differentially expressed if they exhibited absolute log2 fold change ≥ 0.5 and Benjamini–Hochberg adjusted P value < 0.01 within a timepoint. Significantly upregulated genes were further classified based on temporal expression dynamics as transient (significant at one day only), persistent (significant at both one day and ten days), or late (significant at ten days only).

### Gene Ontology analysis

Gene Ontology (GO) enrichment analysis was performed using gProfiler2 in R (version 0.2.3), with a custom background of expressed genes based on expression-filtered RNA-seq genes and false discovery rate (FDR) < 0.05.

### snATAC-seq preliminary analysis

We used SnapATAC2 (2.5.0)^61^ to process fragment files generated from Cell Ranger ARC to make the cell-by-peak h5ad file and bigwig files for each cell type as assigned by their RNA modality clustering. We used the PoissonVI model^62^ from the scVI-tools^63^ to learn a cell-by-peak based latent embedding. Cell level Differential peak accessibility analysis was performed for our cell types of interest (DG, CA3, CA1, and MGE-GABA – which combines SST and PV neurons) within the PoissonVI framework using peaks accessible in >1% of all cells. Significant differentially accessible peaks based on the PoissonVI model were defined as FDR < 0.05, abs(mean log_2_(fold change)) > 0.3 and abs(empirical log_2_(fold change)) > 0.3.

### DG pseudobulked ATAC analysis

Peaks from the Cell Ranger ARC–derived cell-by-peak matrix were retained for downstream pseudobulk analysis if they were accessible (nonzero counts) in at least 2% of all DG annotated cells within either the TRAP+ or TRAP− population across the entire dataset, thereby defining a superset of 104,480 peaks present in at least one condition. Raw counts for these peaks were summed across all DG cells within each sample to generate pseudobulk accessibility profiles. Differential accessibility was assessed on these peaks independently within each timepoint using DESeq2 (v1.46.0), with TRAP status (TRAP+ versus TRAP−) as the variable of interest and replicate included as a covariate to control for batch effects. Statistical significance was assessed using Wald tests with Benjamini–Hochberg correction for multiple testing. Peaks with increased accessibility in TRAP+ relative to TRAP− samples (log2 fold change > 0, adjusted P value < 0.05) were classified based on temporal dynamics as transient (significant at one day only), semitransient (significant at one and two days but not ten days), or persistent (significant at all three timepoints). Heatmaps were generated in Python using matplotlib (version 3.10.6). BigWig files for DG pseudobulk ATAC-seq signal were generated using SnapATAC2 (v2.4) with a bin size of 32. Replicate-level tracks were averaged across biological replicates within each timepoint and TRAP condition using wiggletools (v1.2.3.) to generate consensus tracks for visualization.

### Transcription factor motif enrichment analysis

Motif enrichment analysis was performed using HOMER (findMotifsGenome.pl) on 200 bp windows centered on each peak. Background peak sets for each comparison were generated by selecting peaks with matched GC content and accessibility distributions relative to the corresponding foreground set. For timepoint-specific analyses, foreground peaks were defined as those exhibiting significantly increased accessibility in TRAP+ relative to TRAP− neurons (log2 fold change > 0 and adjusted P value < 0.05), and matched background peaks were selected from the pool of all non-differentially accessible regions (adjusted P value ≥ 0.2 and |log2 fold change| ≤ 0.2 across all timepoints). For comparisons of persistent versus transient TRAP+ enriched differentially accessible regions (DARs), persistent peaks were used as the foreground and a matched set of transient peaks was selected as the background. GC content was computed from mm10 genomic sequences using the bolero package. To quantify AP-1 motif occurrence across the full 500bp peaks, all DG pseudobulk peaks were additionally annotated using HOMER (annotatePeaks.pl), and the proportion of peaks within each DAR kinetic class containing zero, one, or multiple AP-1 motifs was calculated.

### Peak to gene correlation analysis

For peak-to-gene linkage analysis, fragment files generated by Cell Ranger ARC were processed using ArchR (v1.0.3) to generate Arrow files using only cell barcodes corresponding to dentate gyrus (DG) neurons. An initial inclusive peak set was defined using peaks detected in >1% of dentate gyrus (DG) cells in either the TRAP+ or TRAP− population and used to construct the ArchR cell-by-peak accessibility matrix for dimensionality reduction and clustering. This approach maximized feature representation to improve separation of cell states in latent space. Latent semantic indexing (LSI) was performed using default parameters to generate a low-dimensional embedding for downstream cell grouping and pseudocell construction. For downstream analyses, the more stringently defined peak set corresponding to those used in differential accessibility testing (n = 104,480) was used for final peak-to-gene linkage analysis. Gene expression data were incorporated by aligning Cell Ranger ARC–derived raw gene expression matrices to the ArchR project using shared cell barcodes, To control for differences in cell number between conditions, TRAP− cells were randomly downsampled within each replicate and timepoint to match the number of TRAP+ cells. Peak-to-gene links were computed using ArchR (addPeak2GeneLinks) with default parameters (k = 100, knn = 500, overlapCutoff = 0.8), testing correlations between peak accessibility and gene expression within ±500 kb of each gene transcription start site. Significant links were defined as those with correlation > 0.45 (corresponding to Padj values < 1 × 10⁻²³). Significant peak-to-gene links were intersected with sets of significantly upregulated differentially accessible regions (DARs) and differentially expressed genes (DEGs) to identify regulatory relationships between chromatin accessibility and gene expression.

### Cell Level Quantification of AP-1 Motif Accessibility State with Chromvar

For ChromVAR analysis,^64^ we used the whole cell-by-peak matrix as input and scanned motifs across the JASPAR2024 motif database.^65^ We then performed a two-sided t-test with BenjaminiHochberg FDR correction on the cell-level ChromVAR scores and selected significant differential motifs based on an FDR < 0.05. Similar motifs belonging to the same motif family were merged, and the most differential motif was selected to represent the motif family (e.g. FOS for the AP-1 family). Computing distributions of AP-1 chromVAR scores at each timepoint defined upper and lower quartile thresholds in 10d TRAP+ DG cells. These thresholds were subsequently applied to the 1d timepoint cells to identify matched AP-1 chromVAR states across time. Cell barcodes corresponding to these chromVAR-defined subsets were retained for downstream analyses.

#### Chromvar informed peak analysis

To visualize aggregate chromatin accessibility across chromVAR-defined cell subsets, fragment files were filtered by barcode to retain fragments corresponding to each selected cell population. These subset-specific fragment sets were converted to BAM files and merged across biological replicates within each condition. Genome-wide signal tracks were generated using deepTools (bamCoverage) with RPGC normalization. Aggregate accessibility profiles centered on peak regions of interest were computed using deepTools (computeMatrix and plotProfile) using a ±1 kb window around peak centers. Aggregated accessibility profiles were generated for peak sets defined from prior analyses, including peaks linked to differentially expressed genes by Archr as well as transient and persistent DAR classes more generally.

#### Chromvar informed gene expression analysis

To assess the impact of AP-1 chromVAR scores on pseudobulk gene expression profiles of TRAP+ cells, TRAP+ cells were aggregated into subsets based on chromVAR-defined quartiles and compared to matched TRAP− populations using a bootstrapped differential expression framework. For each comparison, TRAP+ subsets (e.g., Q1 or Q4) were compared to a shared set of 100 randomly sampled TRAP− populations, selected independently of chromVAR score and approximately matched in size, ensuring that effect sizes were evaluated relative to identical baseline populations across subsets. Genes with adjusted P value < 0.05 and log2 fold change > 0 in more than 50% of iterations were defined as robustly detected differentially expressed genes (DEGs), and the median log2 fold change across iterations was used as the effect size. The union of upregulated DEGs identified in either 10d Q1 versus TRAP− or 10d Q4 versus TRAP− comparisons was taken forward for comparison of effect sizes between these subsets. The linkedpoint boxplot of median Log2 Fold Change values shown in was generated by comparing how the magnitude of gene expression differences relative to TRAP− cells varied across subsets, enabling assessment of each subset’s relative contribution to the 10-day ensemble gene expression program. Direct comparison between Q1 and Q4 TRAP+ subsets for the supplementary volcano plot was performed using a single DESeq2 analysis restricted to these populations.

### Quantification of cell-level transcriptional heterogeneity (Transcriptional Program Score)

To quantify heterogeneity in the engagement of the ensemble-associated transcriptional program at the single-cell level, a gene set–based scoring approach was used. The top 100 genes upregulated at one day post-conditioning (ranked by Padj from pseudobulk differential expression analysis) were selected to define an ensemble gene program. The resulting transcriptional program scores (TPS) were computed for each cell using Scanpy (sc.tl.score_genes), yielding a program activity score per cell. Distributions of program scores were compared across TRAP− cells and TRAP+ cells at each timepoint using density plots. To define a threshold for robust program activation, a baseline was established using TRAP− cells by computing the 90th percentile of program scores within each replicate and timepoint, and averaging these values to obtain a global threshold. Cells with program scores exceeding this threshold were classified as transcriptionally active with respect to the ensemble gene program (TPS-high). The proportion of TPS-high cells was then calculated for each condition and timepoint, enabling quantification of the fraction of TRAP+ neurons exhibiting sustained transcriptional engagement relative to TRAP− cells and assessment of temporal changes in transcriptional heterogeneity. To assess the relationship between AP-1 activity and engagement of the ensemble-associated transcriptional program at the single-cell level, AP-1 (FOS) chromVAR deviation scores were directly correlated with RNA program scores across cells, and significance was evaluated using Spearman correlation.

### Spatial Distribution Matching

Cells were integrated with a BICCN MERFISH reference atlas2 using transcriptional nearestneighbor mapping, and spatial coordinates (x, y, z) were imputed by transferring the coordinates of matched reference cells. Deviations in spatial makeup of the TRAP+ and TRAP-populations were observed in the Y (dorsal-ventral) coordinate axis in CA3 in particular. To control for spatial confounding, cells were stratified by subclass, timepoint, and replicate, and Trap and Neg populations were distribution-matched along the dorsal–ventral axis by binning and subsampling to equalize spatial densities while retaining maximal cell numbers. Differential expression analysis was performed as described above.

### Electrophysiology

#### Acute hippocampal slice preparation

Acute coronal hippocampal slices were prepared from mice either 2 or 9/10 days following contextual fear conditioning (postnatal day 39-46). Mice were anesthetized with isoflurane and perfused with an ice-cold N-methyl-D-glucamine (NMDG)-based artificial cerebrospinal fluid (NMDG-ACSF) equilibrated with 95% O_2_/5% CO_2_ consisting of (in mM): 93 NMDG, 2.5 KCl, 1.25 NaHPO_4_, 30 NaHCO_3_, 20 Hepes, 25 Glucose, 5 Na-Ascorbate, 3 Na-Pyruvate, 0.5 CaCl_2_, 10 MgCl_2_ (pH adjusted to 7.4). Subsequently, the brain was rapidly removed from the skull and placed in ice-cold NMDG-ACSF. Hippocampal slices (300 µm) were collected with a vibratome (Leica VT1200S). Slices were incubated for 12 minutes at 32°C in NMDG-ACSF, then transferred to room temperature ACSF equilibrated with 95% O_2_/5% CO_2_ consisting of (in mM): 127 NaCl, 25 NaHCO_3_, 1.25 NaH_2_PO_4_, 2.5 KCl, 1 MgCl_2_, 10 glucose and 2 CaCl_2_. Slices were incubated for 1 hr in room temperature ACSF before being transferred to the recording chamber.

#### Whole-cell Patch-Clamp Electrophysiology

Whole-cell patch-clamp electrophysiology was performed using an upright Olympus BX51 WI microscope with a 60X water immersion objective (Olympus Lumplan FI/IR 60x/0.90 numerical aperture), equipped with an Axon Instruments CV-7B head stage, an Axon MultiClamp 700B amplifier, and an Axon Digidata 1440A digitizer. For visualization, neurons were imaged using either an Andor Zyla sCMOS (ZYLA-5.5) camera or an infrared CCD (Dage-MTI IR-1000) camera. Epifluorescence was illuminated with an Excelitas X-Cite 120LED system to distinguish trapped from non-trapped cells. All recordings were conducted in the granule cell layer of the medialventral suprapyramidal blade of the dentate gyrus. All non-trapped cells recorded were located within the neighboring vicinity of trapped cells.

All recordings were performed with 3-6 MΩ borosilicate glass with filament patch pipettes (Sutter BF150-86-7.5). Voltage-clamp recordings were filtered with a 1 kHz Bessel filter and sampled at 100 µs. Current-clamp recordings were filtered with a 10 kHz Bessel filter and sampled at 100 µs. Whole-cell patch-clamp electrophysiology was performed in ACSF equilibrated with 95% O_2_/5% CO_2_ at room temperature. To isolate miniature excitatory and inhibitory post-synaptic currents Tetrodotoxin citrate (TTX) (0.5 µM) and D-AP5 (50 µM) were included in the bath solution. Miniature excitatory post-synaptic currents were recorded at –70 mV, and inhibitory miniature postsynaptic currents were recorded at 0 mV.

For voltage-clamp recordings, internal pipette solution contained (in mM): 127 CsMeSO_3_, 10 CsCl, 10 HEPES, 0.5 EGTA, 2 MgCl_2_, 0.16 CaCl_2_, 2 MgATP, 0.4 NaGTP, 14 Na_2_-phosphocreatinine, and 2 QX314-Cl (pH 7.2, 295 mOsm). For current clamp recordings, internal pipette solution contained (in mM): 142 K^+^-gluconate, 4 KCl, 10 HEPES, 4 MgATP, 0.3 NaGTP, 10 Na_2_phosphocreatinine, and 1.1 EGTA (pH 7.2, 280 mOsm).

#### Data Acquisition and Analysis

Clampex 10.6 or 10.7 software (Molecular Devices) was used for data acquisition. Mini Analysis software (Synaptosoft) was used to calculate miniature post-synaptic current frequencies, amplitudes, and kinetics. The open-source software Stimfit^66^ was used to analyze intrinsic properties. Series resistance was monitored throughout each recording and if it was changing for more than 20% during the course of the recording or was > 30 MΩ the recording was not included. All analyses were conducted blind to cell identity. GraphPad Prism 10 was used to conduct all statistical analyses. A D’Agostino & Pearson test for normality was run to select the appropriate parametric or nonparametric statistical analysis to compare groups. p< 0.05 was considered statistically significant. Sample sizes were not pre-determined but are comparable to those used in previous studies reporting similar datasets^9,60,67^.

### CUT&RUN, library preparation and sequencing

Juvenile, four-six week old, WT or FFJ^flox^ mice injected with cre-recombinase at p33-p37 were injected with saline or Kainic acid (16 mg/kg, Sigma K0250) and tissue was isolated either 2 hours, 1 or 5 days post injection. Mice were deeply anesthetized with isoflurane, decapitated and brains were placed in a stainless-steel brain matrix on ice to cut 1 mm sections of anterior to posterior hippocampus. Sections were transferred into ice-cold PBS and the dentate gyrus was isolated under a dissection microscope. Nuclei were prepared as described for FANS sorting for Multiome experiments and NeuN+ nuclei, as well as NeuN+/GFP+ nuclei for FFJ^cKO^ were sorted into 2%

Homogenization Buffer by FANS. Nuclei from two mice were pooled for each biological replicate, and ∼600.000 neuronal nuclei were isolated for each biological replicate. A total of four biological replicates for each time point were collected across two independent batches.

CUT&RUN experiments were performed essentially as previously described^68,69^. Nuclei were pelleted by centrifugation at 500xg for 10 min at 4C, then resuspended in wash buffer (20mM HEPES pH 7.5, 150mM NaCl, 0.2% Tween-20, 0.1% BSA, 0.5mM spermidine, supplemented with protease inhibitors). Magnetic Concanavalin A beads (Bangs Laboratories BP531) were activated twice with binding buffer (20 mM HEPES-KOH pH=7.9, 10mM KCl, 1mM CaCl_2_, 1mM MnCl_2_), then added to the nuclei. Nuclei were incubated with beads for 10min at room temperature. The bead-bound nuclei were resuspended in antibody buffer (20mM HEPES pH 7.5, 150mM NaCl, 0.2% Tween-20, 0.1% BSA, 0.1% Triton X-100, 2mM EDTA, 0.5mM spermidine, supplemented with protease inhibitors), and each sample was divided into three reactions of 200,000 nuclei in 50µl each. Primary antibody was added and incubated at 4°C overnight with gentle rocking. Primary antibodies used were rabbit anti-FOS (in-house 1096AE, 1:50)^60^, rabbit anti-FosB (Cell Signaling 2251, clone 5G4, lot 5, 1:50), rabbit IgG isotype control (Cell Signaling 2729, lot 11, 1:50). Nuclei were washed with antibody buffer, then incubated with protein Aconjugated MNase (700ng/ml in antibody buffer, in house) for 1 hr at 4C. Nuclei were then washed twice with Triton wash buffer (20mM HEPES pH 7.5, 150mM NaCl, 0.2% Tween-20, 0.1% BSA, 0.1% Triton X-100, 0.5mM spermidine, supplemented with protease inhibitors). FOSB samples were washed once with low-salt wash buffer (20mM HEPES pH 7.5, 0.2% Tween-20, 0.1% Triton X-100, 0.5mM spermidine, supplemented with protease inhibitors) and then incubated in calcium incubation buffer (FOS, IgG, Triton wash buffer with 3mM CaCl_2_) or low-salt calcium incubation buffer (FOSB, 3.5mM HEPES pH 7.5, 10mM CaCl2, 0.2% Tween-20, 0.1% Triton X-100, supplemented with protease inhibitors) for 45 min in a wet ice bath. The reaction was terminated by adding 2x STOP buffer (340mM NaCl, 20mM EDTA, 4mM EGTA, 0.04% Triton X-100, 10pg/ml yeast spike-in DNA and 0.1mg/ml RNAse A) and samples were incubated at 37°C for 20 min to release fragments from chromatin. Samples were then centrifuged at 16,000g for 5min at 4°C. Supernatant was transferred into a fresh tube with 0.1% SDS and 0.2mg/ml proteinase K and incubated at 65°C for 1 hr with 500 rpm shaking. Fragments were purified with standard phenolchloroform-isoamyl alcohol extraction and ethanol precipitation.

CUT&RUN libraries were prepared using the ThruPLEX DNA-seq kit (Takara R400676) following standard protocol with the following modifications: Library amplification PCR cycling was performed as at 72°C for 3 minutes, 85°C for 2 minutes, 98°C for 2 minutes and 4 cycles of [98°C for 2 seconds, 67°C for 20 seconds, 72°C for 30 seconds], 9 cycles of [98°C for 20 seconds, 72°C for 15 seconds], 4°C hold. Libraries were purified with a 0.5x-1.7x AMPure beads (Beckman Coulter A63881) clean up. Libraries were sequenced with paired ends on an Illumina NextSeq 500 using a 75-cycle high output kit.

### CUT&RUN Data preprocessing and analysis

Reads were trimmed for adapters and quality using cutadapt (5.1), then mapped to mm10 using bowtie2 (2.5.4) with the following settings: bowtie2 –-local –-very-sensitive-local –-dovetail –-nounal--no-mixed –-no-discordant –-phred33. Mapped reads were sorted and indexed using samtools (1.21), de-duplicated using picard (3.4.0) MarkDuplicates, and filtered to <=120 bp fragment size using alignmentSieve (deepTools 3.5.6). For visualization of individual genomic loci, CPM-normalized bigwig coverage files were generated using bamCoverage (deepTools 3.5.6) and visualized in IGV. For depiction of CUT&RUN signal over regions of interest previously defined from the scATAC-seq, fragment size-filtered reads were counted, normalized to sequencing depth, and visualized using DiffBind (3.16.0, in R 4.4.0).

### Histology and Fluorescent in-situ hybridization (FISH)

Mice were deeply anesthetized with 90 mg/kg ketamine and 50 mg/kg xylazine by intraperitoneal injection, and transcardially perfused with ice-cold PBS followed by 4% paraformaldehyde (PFA) in PBS. Dissected brains were post-fixed overnight at 4°C, incubated overnight in 15% sucrose prepared in PBS, and then overnight in 30% sucrose in PBS at 4°C. Brains were sectioned on a vibratome LeicaVT1000 at 30 µm, cryoprotected in 1.5 parts ethylene glycol, 1.5 parts glycerol and 2 parts 1X PBS and frozen in cryoprotective 1.5ml tubes at –80°C until further use. Before further processing, free-floating hippocampal sections were washed in 1X PBS three times, permeabilized with 0.25% Triton X-100 in 1X PBS for 1 h, and blocked (5% bovine serum albumin, 0.3% Triton X-100, and 10% donkey serum in 1X PBS) for 2 h. Blocking solution was replaced by primary antibody solution (1% BSA, 0.3% Triton X-100 and 5% donkey serum in 1X PBS). Sections were incubated overnight at 4°C with the following primary antibodies: c-Fos anti-guinea pig (1:3000, Synaptic Systems, #226308) and FosB anti-rabbit (1:1000, Cell Signaling Technology #2251). Endogenous mcherry signal in Trap+ neurons was amplified using goat (1:2000, antibodies-online # ABIN1440058). Brain slices were subsequently washed three times in 0.1% Triton 100X in PBS for 15 min each. Sections were incubated in secondary antibody solution (1% BSA, 0.3% Triton X-100 and 5% donkey serum in PBS 1X) for 2 hr at room temperature with the following secondary antibodies: Cy3 donkey anti-rabbit IgG (1:1000, Jackson ImmunoResearch, #711-165-152) and Cy5 donkey anti-guinea pig IgG (1:1000, Jackson ImmunoResearch, #706175-148), Cy3 donkey anti-goat (1:1000, Jackson ImmunoResearch #705-165-147 or Cy5 donkey anti-rabbit IgG (1:1000, Jackson ImmunoResearch #711-175-152). Slices were washed twice at room temperature with 0.1% Triton 100X in PBS and then counterstained with 4’,6diamidine-2’-phenylindole dihydrochloride (DAPI) (1:5000, Sigma Millipore, #MBD0015) in PBS for 5 min. Finally, brain slices were washed twice in 1X PBS for 15min and mounted using ProLong Diamond Gold Antifade (Life Technologies #P36970).

For Fluorescent-in-situ hybridization (FISH) all solutions were prepared in RNase-free PBS and water. Sections were prepared and frozen as described above. After thawing and washing in 1X PBS, sections were mounted on RNase-free SuperFrost Plus slides (ThermoFisher) and probed against genes of interest using the RNAscope Multiplex Fluorescent Assay v2 protocol (ACDBio, Catalogue #323110). RNAscope, was performed following manufacturer’s instructions, with the exception of hydrogen peroxide solution from the kit which was replaced with 30% hydrogen peroxide solution in water (Sigma Aldrich, Catalogue #216763), and Protease IV was used for 19 min. The following probes were used: Mm-Sorcs3-C1 (#473421) and Mm-Fos-C2 (316921-C2). Protein signal for mcherry and FosB (using same antibodies as described above) in Trap+ neurons was performed on slide using histology procedures described above. As a final step, sections were counterstained with 4’,6-diamidine-2’-phenylindole dihydrochloride (DAPI) (1:5000, Sigma Millipore, # MBD0015) in PBS for 5 minutes, washed in PBS 1X and mounted using ProLong Diamond Gold Antifade. Images were acquired using z-stacks of 30 µm step size with 40X objectives, at 1024 x 1024 pixel resolution, and 16-bit depth on a ZEISS LSM800 confocal microscope.

### Nuclear segmentation, FISH and immunostaining analysis

Following imaging, raw.czi files iteratively underwent nuclear (DAPI+) and cellular (TRAP+) segmentation in three-dimensional space using a custom trained model of the Cellpose-Segment Anything Model (CPSAM) in Cellpose (v4.0.4) using python (3.10.13) and torch (2.6.0+cu124!) run in GPU-enabled mode on the Harvard Medical School O2 HPC cluster. DAPI+ nuclei (diameter = 30 pixels) and TRAP+ cells (diameter = 100 pixels) were segmented separately after adjusting for voxel size. Next, DAPI+ and TRAP+ masks were overlaid, and all DAPI+ and TRAP+ masks were compared for greatest overlap in 3-D space. When any single DAPI+ and TRAP+ mask were found to mutually overlap the most of all other masks and the DAPI+ mask overlapped at least 70% in 3-D space with its mutually co-assigned TRAP+ mask, this DAPI+/TRAP+ nucleus was considered paired and used for subsequent TRAP+ analysis for FISH and protein stain quantification. To limit the possibility of signal bleed-through in the Z axis, all TRAP-nuclei directly neighboring a TRAP+ nucleus were removed from downstream analysis. To diminish the effect of poorly segmented or doublet nuclei, segmented nuclear volumes were filtered to be at least 7500 voxels large. After segmentation, we manually performed quality control curation of all TRAP+ nuclei by first forming a maximum projection Z-stack for each protein or FISH signal channel limited to the 3-D region defined by the DAPI+/TRAP+ mask boundaries per cell. Rarely, DAPI+ or TRAP+ masks were found to be poorly paired or significant spectral bleed-through was noted and these individual nuclei were removed from downstream analysis. After quality control, raw intensities without any background subtraction of FISH and protein signal in TRAP– or TRAP+ nuclear masks were considered for quantification. The sum of this signal present per channel across the each DAPI+ mask was normalized to its nuclear volume and further normalized against the animal-specific median intensity value across all cells measured for that animal. These median-normalized volumetric intensities were used for comparison. Positivity thresholds for the Fos RNA, FOS Protein, and FOSB protein signals were computed through manual evaluation of the per cell maximum intensity projections to identify correspondence between per-animal median normalized volumetric intensity values and positivity status as classified separately by two experimenters. These cutoffs were then applied against additional nuclei masks and checked post hoc to evaluate cutoff accuracy. This process was repeated until a cutoff was selected with >90% accuracy in over 100 more manually evaluated cells. Sorcs3 intensity bins were determined on per animal basis by ranking the TRAP+ cells in terms of their volumetric Sorcs3 RNA intensity and selecting tertiles as cutoff points for Low (bottom tertile), Mid (middle tertile), and High (top tertile) Sorcs3 expression levels.

### Statistical Testing for Imaging Datasets

Comparisons between TRAP+ and TRAP− populations were performed using paired t-tests on animal-level summary statistics within each timepoint. Summary values were calculated per animal as either the median normalized intensity across all cells of each TRAP status (for continuous measures) or the percentage of cells positive for signal (for binary readouts). For comparisons across timepoints or between recall and non-recall conditions, analysis of variance (ANOVA) was performed with TRAP status treated as a within-animal factor and timepoint or recall condition as between-animal factors. Multiple testing correction was applied using the Benjamini–

Hochberg method. To assess the distribution of TRAP+/Fos+ cells across Sorcs3 RNA intensity tertiles or the distribution of TRAP+/Sorcs3-High cells across FOSB protein intensity tertiles, we tested deviation from a null expectation of equal occupancy (1/3 per tertile). The proportion of the target population within each tertile was calculated, and a parametric approach was taken to evaluate whether observed distributions differed from the expected uniform distribution while preserving per-animal totals. Deviation from a uniform distribution was assessed using a Monte Carlo goodness-of-fit test to derive empirical p-values from 100,000 simulations.

### Code availability

All code used in this study was previously published and is cited in the Methods section.

## Acknowledgements

We thank members of the Greenberg laboratory, Gord Fishell, and Sandeep Robert Datta for helpful discussions over the course of this project and on the manuscript, as well as Peter Scheiffele and Christopher D. Harvey for feedback on the manuscript. We would like to especially thank Erin E. Duffy for preparing the Rshiny database. We would further like to thank the Boston Children’s Hospital Flow Cytometry and viral core facilities. The Greenberg laboratory is supported by the Tan-Yang Autism Center at Harvard Medical School and we thank Steve and Judy Kaye for their generous support of this research during a difficult time. S.K. was supported by a Y. Eva Tan Fellowship of the Tan Yang Autism Center, N.J.G. was supported by Developmental Neurology Institutional Training Program T32007473, H.L. was supported by the Harvard Society of Fellows, S.A.K. was supported by the National Science Foundation Graduate Research Fellowship Program (DGE 2140743), L.X. was supported by NIH/NCI T32CA272386 and a Lefler Fellowship from the Edward R. and Anne G. Lefler Center, M.E.G. was supported by funding from the NIH and an R35NS143029. L.T. was supported by an EMBO Postdoctoral Fellowship (ALTF 167-2020), Long-term Human Frontiers Science Program Fellowship (LT000340) and the Charles A. King Trust Postdoctoral Research Fellowship. The funders had no role in study design, data collection and analysis, or the decision to prepare and publish the manuscript.

## Author contributions

I.R.V., M.E.G. and L.T. conceived and designed the study. I.R.V. and L.T. designed, executed and analyzed the majority of the experiments, including collection of all of the provided datasets. In addition, I.R.V. performed behavioral experiments and bioinformatic, S.K. provided intellectual guidance in experimental design and contributed to data collection, N.J.G. and L.T. performed electrophysiology experiments, H.L. assisted with bioinformatic analysis, S.K.A. and I.R.V. performed analysis of imaging data, L.X. performed CUT&RUN and analysis, S.S. provided technical assistance for FISH and image acquisition, E.C.G. provided intellectual guidance for design of the study and writing of the manuscript. I.R.V., M.E.G. and L.T. wrote the manuscript with input from all authors.

## Data availability

The snMultiome sequencing and CUT&RUN data generated have been deposited in the Gene Expression Omnibus database and will be made available upon publication of the manuscript. A web-based searchable database, where researchers can look up genes of interest and extract plots of gene expression across major cell classes in TRAP+ versus TRAP-neurons will be made available from our lab website upon publication.

## Supplementary Materials

**Supplementary Table 1**: Numbers of nuclei per cell class and replicate and DESeq2 normalized count matrix for genes detected in each replicate.

**Supplementary Table 2**: Pseudobulk differential gene expression analysis for all five cell classes analyzed

**Supplementary Table 3**: Pseudobulk analysis of differential peak accessibility for DG, cell-bycell level analysis of accessible chromatin regions, ArchR-linked peaks to genes and TPS

**Supplementary Table 4:** Analysis of Fluorescent in-situ hybridization

**Supplementary Table 5**: Pseudobulk differential gene expression analysis for all five cell classes analyzed that are corrected for spatial origin of nuclei.

