## Supplementary Figures for "Long-lasting chromatin and transcriptional states shape the maintenance of memory ensembles"

### Figure S1

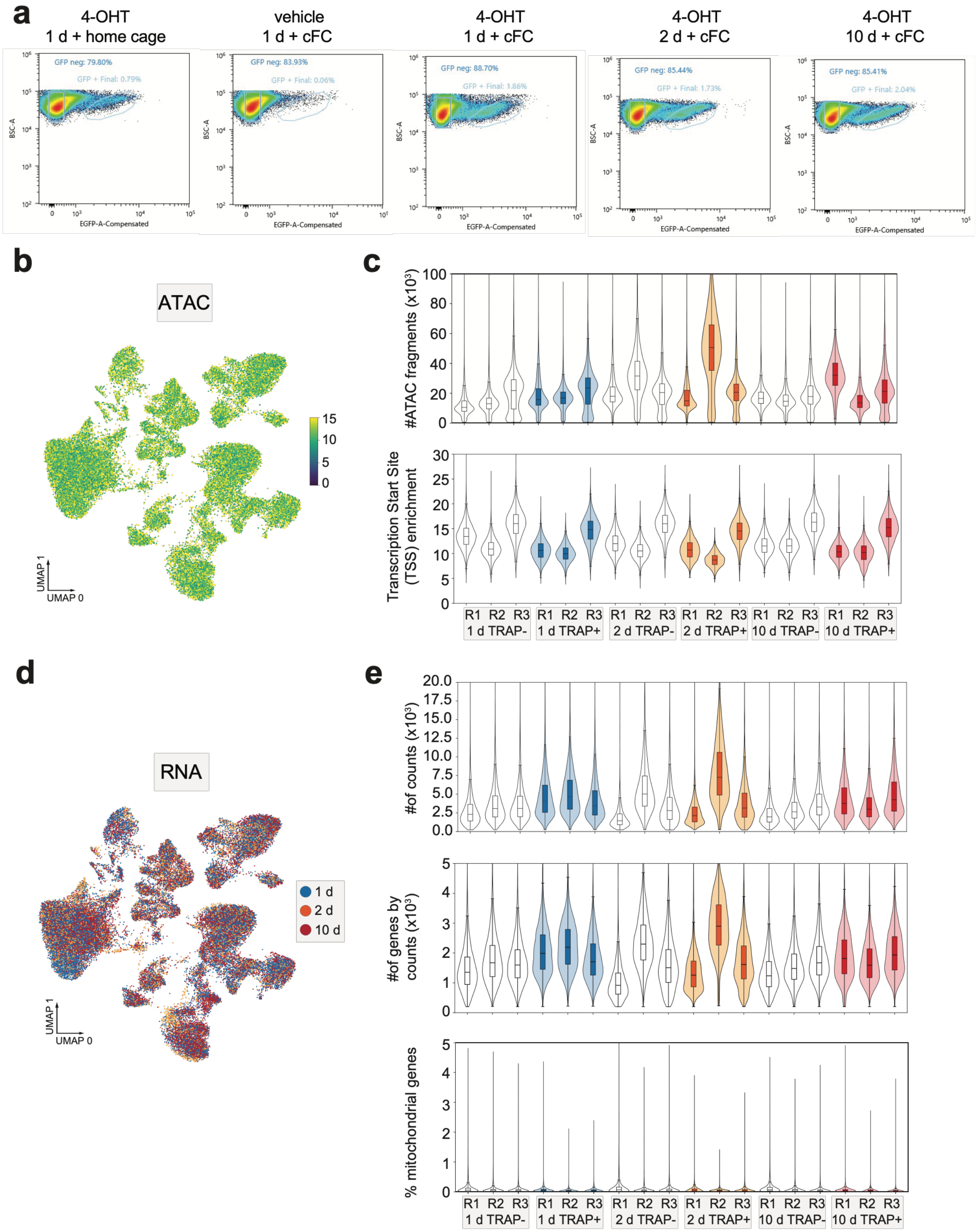
a
4-OHT
1 d + home cage
vehicle
1 d + cFC
4-OHT
1 d + cFC
4-OHT
2 d + cFC
4-OHT
10 d + cFC
GFP neg: 79.80%
GFP + Final: 0.79%
GFP neg: 83.93%
GFP + Final: 0.06%
GFP neg: 88.70%
GFP + Final: 1.86%
GFP neg: 85.44%
GFP + Final: 1.73%
GFP neg: 85.41%
GFP + Final: 2.04%
BSC-A
EGFP-A-Compensated
b
ATAC
UMAP 1
UMAP 0
c
#ATAC fragments (x10³)
Transcription Start Site (TSS) enrichment
R1 R2 R3
1 d TRAP-
1 d TRAP+
2 d TRAP-
2 d TRAP+
10 d TRAP-
10 d TRAP+
d
RNA
1 d
2 d
10 d
UMAP 1
UMAP 0
e
#of counts (x10³)
#of genes by counts (x10³)
% mitochondrial genes
R1 R2 R3
1 d TRAP-
1 d TRAP+
2 d TRAP-
2 d TRAP+
10 d TRAP-
10 d TRAP+

**Supplementary Figure 1: Single-nucleus multiome dataset quality metrics, related to Figure 1**

a) Representative FANS traces of control (TRAP+ neurons in home cage treated with 4OHT, or post-cFC with vehicle injection) and experimental (TRAP+ neurons at one, two- or ten-days post-cFC when treated with 4OHT) conditions. Percentages of TRAP+ (GFP+) neurons are indicated.
b) UMAP as in Figure 1b for transcription start site enrichment (TSSE) for each cell and each replicate in the dataset.
c) Violin plots of number of ATAC fragments and TSSE by TRAP status, replicate (R) and time point.
d) UMAP as in Figure 1b, color-coded by time point.
e) Violin plots of number of counts per cell, number of genes by counts and percentage of mitochondrial RNA detected per cell separated by TRAP status, replicate (R) and time point.

### Figure S2

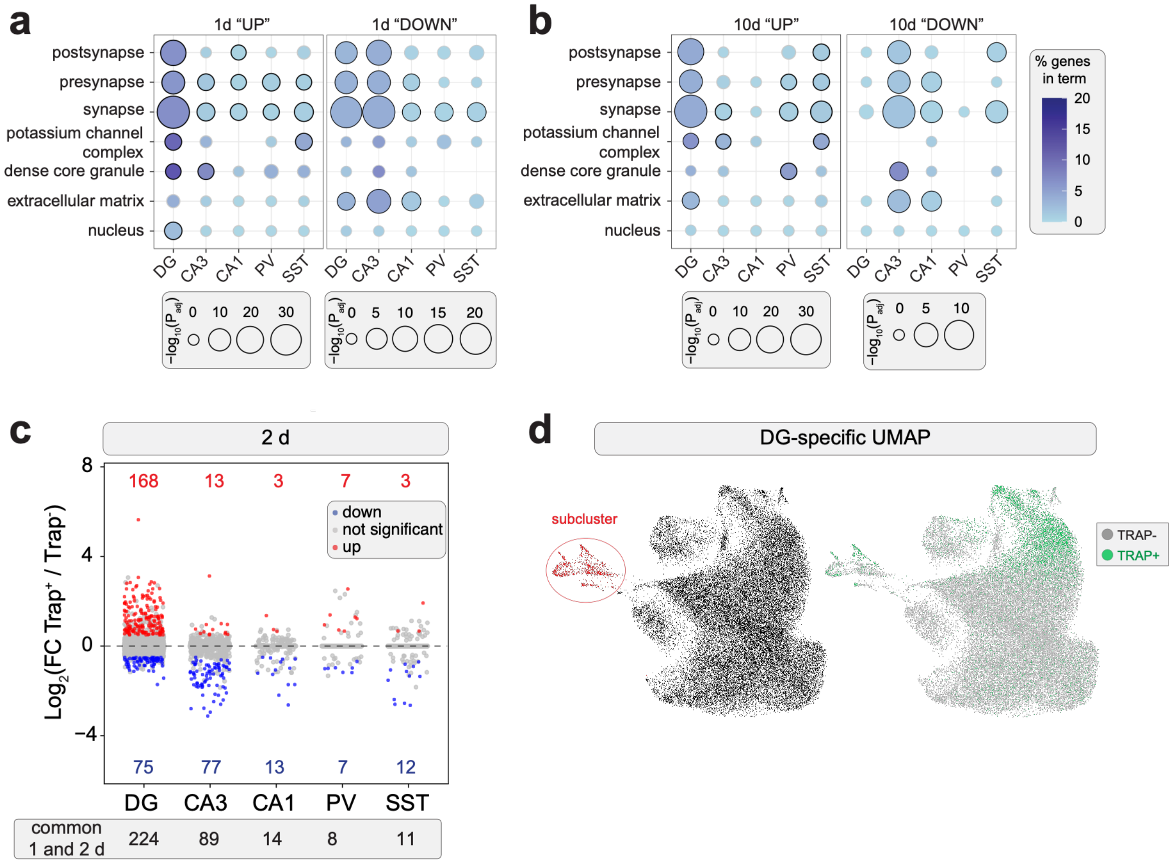


**Supplementary Figure 2: Differential gene expression analysis of TRAP+ and TRAP- neurons over time, related to Figure 1**

a) Dot plots of top enriched Gene Ontology terms of cellular compartment for DG, CA3, CA1, PV and SST up-regulated genes one day post-cFC in Figure 1c (FDR < 0.05, significant terms are outlined in black).
b) Dot plots of top enriched Gene Ontology terms for DG, CA3, CA1, PV and SST up-regulated genes ten days post-cFC in Figure 1c (FDR < 0.05, significant terms are outlined in black).
c) Jitter plots of pseudobulk DESeq2 differential gene expression analysis between TRAP+ and TRAP- neurons within the five major cell classes at two days post-cFC. Genes that are significantly up- or down-regulated in TRAP+ neurons are marked in red or blue, respectively ($Log_2FC \pm 0.5$, adjusted p-value $\leq 0.01$, two-sided Wald test with Benjamini Hochberg correction for multiple testing). Numbers of significantly differentially regulated genes that are either up- or down-regulated are indicated. Numbers of differential genes commonly affected at one and two days for each cell class are shown below.
d) UMAP of DG neuron clustering illustrates a small cluster (red) that contributes to lower count late-emerging genes at ten days post cFC (left). TRAP status in DG cluster is highlighted (right).

### Figure S3

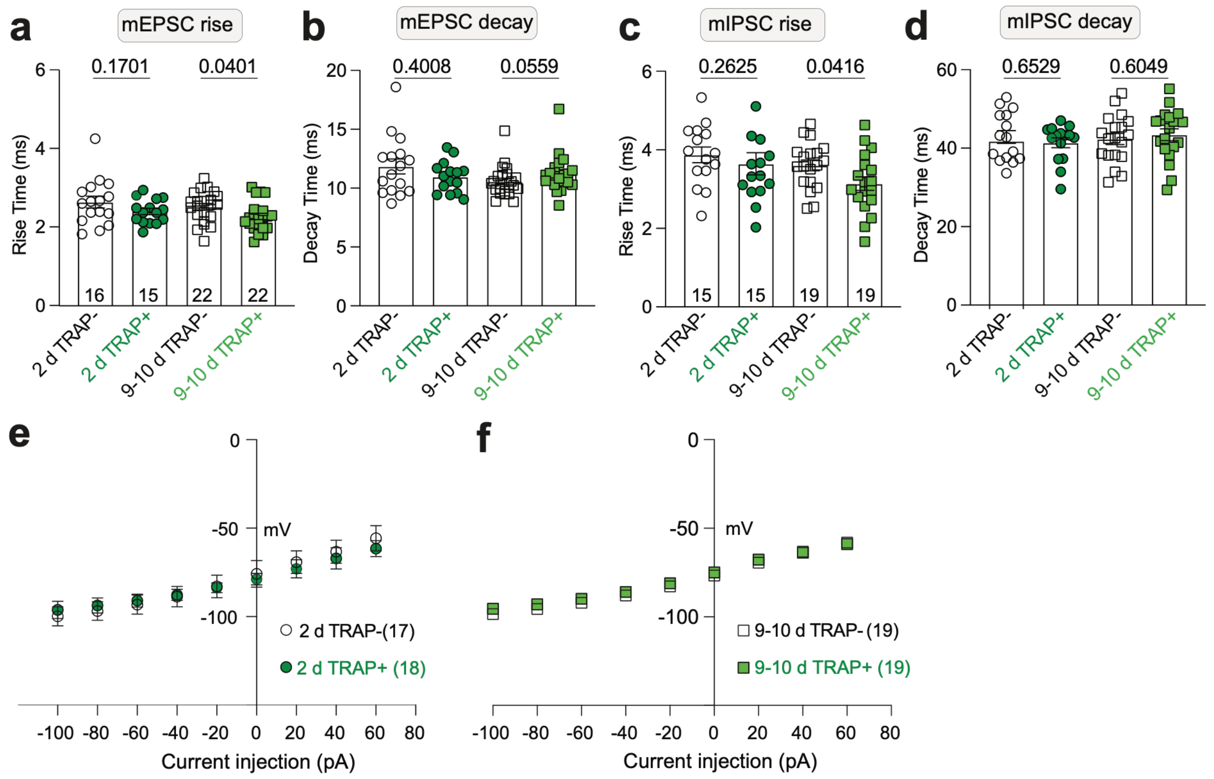


**g**

| | 2 d TRAP- | 2 d TRAP+ | $p\text{-value}_{adj}$ |
|---|---|---|---|
| RMP (mv) | -75.77 ± 1.8 | -78.84 ± 0.78 | 0.1434 |
| Capacitance (pF) | 55.71 ± 3.4 | 54.73 ± 3.6 | 0.8477 |
| AP threshold (mV) | -34.75 ± 1.7 | -34.23 ± 1.2 | 0.8115 |
| AP peak (mV) | 56.81 ± 1.8 | 55.7 ± 2.4 | 0.8802 |
| AP half width (ms) | 1.433 ± 0.06 | 1.425 ± 0.07 | 0.9292 |
| Rise time (ms) | 0.2479 ± 0.01 | 0.2359 ± 0.01 | 0.5102 |
| Max slope decay | 58.70 ± 2.9 | 59.39 ± 3.6 | 0.8102 |
| Latency (ms) | 179.7 ± 4.3 | 177.2 ± 8.0 | 0.3265 |
| n of cells | 16 | 17 | |

**h**

| | 9-10 d TRAP- | 9-10 d TRAP+ | $p\text{-value}_{adj}$ |
|---|---|---|---|
| RMP (mv) | -85.29 ± 1.4 | -78.84 ± 1.2 | 0.2467 |
| Capacitance (pF) | 65.85 ± 4.0 | 70.66 ± 2.9 | 0.3407 |
| AP threshold (mV) | -31.81 ± 1.4 | -33.87 ± 1.4 | 0.4955 |
| AP peak (mV) | 56.74 ± 0.83 | 57.63 ± 1.09 | 0.3224 |
| AP half width (ms) | 1.382 ± 0.06 | 1.333 ± 0.05 | 0.5509 |
| Rise time (ms) | 0.2502 ± 0.01 | 0.2500 ± 0.009 | 0.8350 |
| Max slope decay | 58.75 ± 2.9 | 62.27 ± 2.4 | 0.3635 |
| Latency (ms) | 182 ± 2.7 | 180.1 ± 3.7 | >0.9999 |
| n of cells | 19 | 20 | |

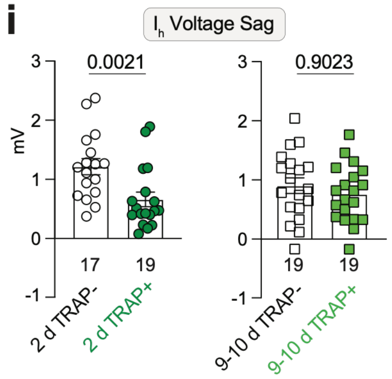

**Supplementary Figure 3: Electrophysiological properties of DG memory ensemble neurons over time, related to Figure 2**

a-d) Miniature excitatory and inhibitory post-synaptic current (mEPCS, mIPSC) rise and decay times in milli-seconds.

e-f) Membrane potential of DG granule cells as a function of injected current.

g-h) Characterization of intrinsic properties.

j) $I_h$ current voltage sag measurements in mV. For all panels, TRAP+ neurons are shown in green, TRAP- neurons in white. Numbers of cells are indicated for each measurement. Numbers of animals as in main Figure 2. Statistics were calculated based on the appropriate unpaired t-test based on normality distribution and standard deviation. Mean ± SEM are shown

### Figure S4

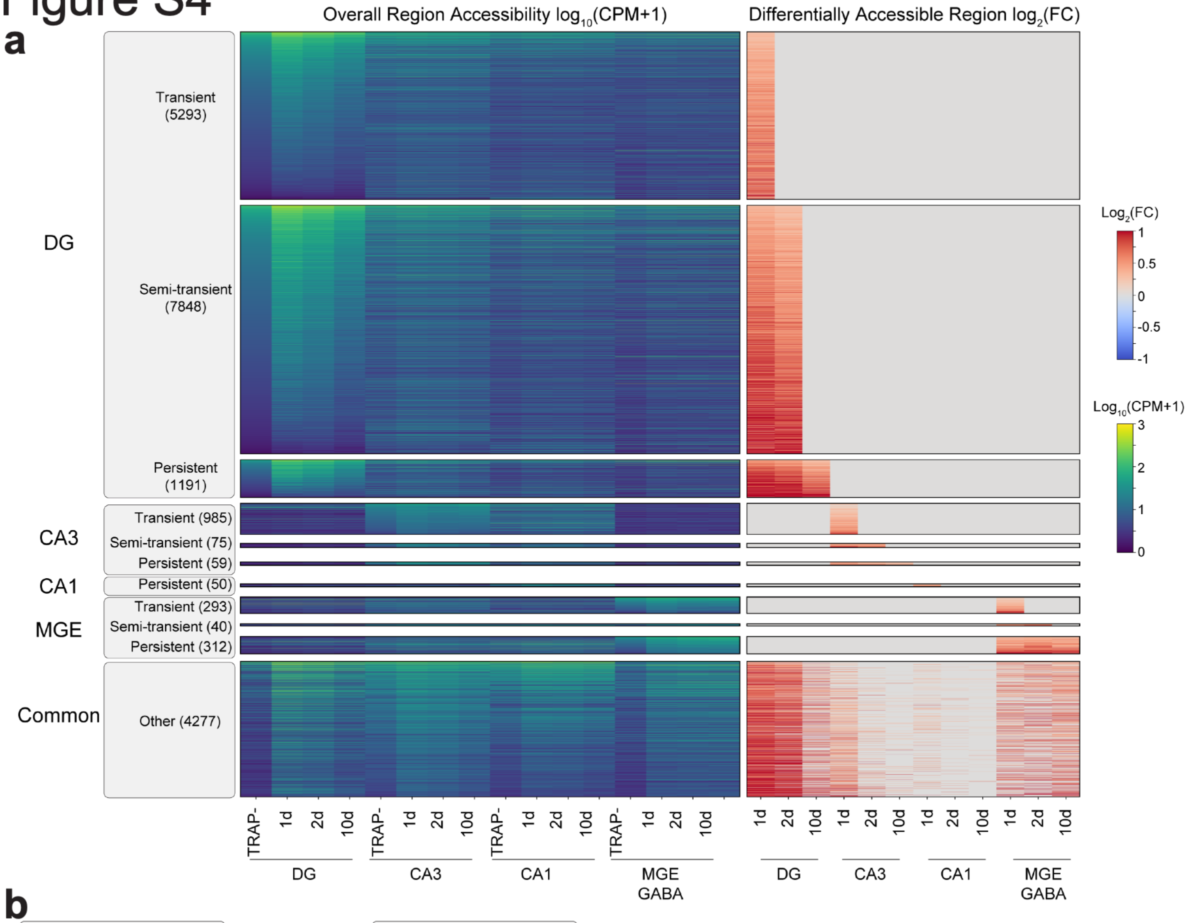


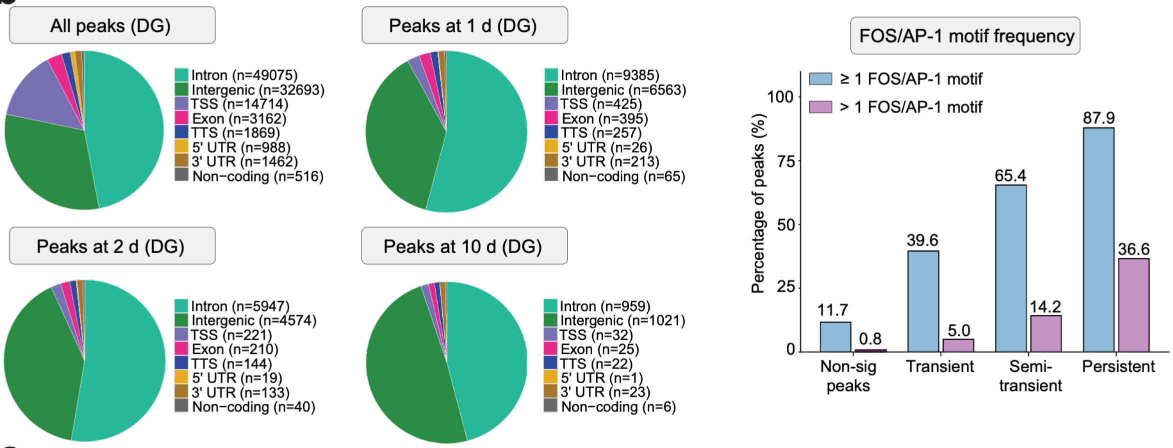


**c**

**1 d**

| Motif (*de novo*) | TF | p-value | target / bkg |
|---|---|---|---|
| 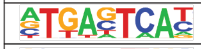 | AP-1 | $1x10^{-4451}$ | 50.57 / 8.62 |
| 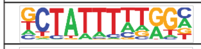 | MEF2 | $1x10^{-76}$ | 14.35 / 9.88 |
| 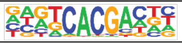 | NPAS4 | $1x10^{-54}$ | 9.89 / 6.72 |
| 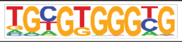 | EGR | $1x10^{-47}$ | 18.60 / 14.58 |
| 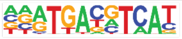 | CREB | $1x10^{-23}$ | 6.48 / 4.77 |

**2 d**

| Motif (*de novo*) | TF | p-value | target / bkg |
|---|---|---|---|
| 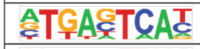 | AP-1 | $1x10^{-4195}$ | 60.91 / 8.68 |
| 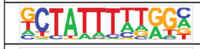 | MEF2 | $1x10^{-51}$ | 15.33 / 10.67 |
| 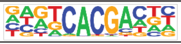 | NPAS4 | $1x10^{-45}$ | 10.39 / 6.77 |
| 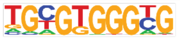 | EGR | $1x10^{-31}$ | 18.12 / 14.31 |
| 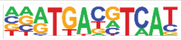 | CREB | $1x10^{-11}$ | 6.08 / 4.63 |

**Supplementary Figure 4: Differential peak accessibility and transcription factor motif analysis of DG memory ensemble neurons, related to Figure 3**

a) Heatmaps of cell-by-cell analysis of differential chromatin accessibility, $\log_{10}$(counts per million) (left) and TRAP-enriched $\log_2$(fold change) (right) across excitatory DG, CA3, CA1 neurons as well as MGE GABA (PV+SST neurons) with distinct transient, semi-transient, persistent and other accessibility kinetics (POISSONVI differential accessibility package-defined significant peaks using Bayesian 'change' test with delta=0.1 and FDR<0.05).
b) Pie charts of genomic location of TRAP+-enriched peaks. Numbers of peaks in each category are indicated.
c) *De novo* HOMER transcription factor motif analysis for the top enriched motifs at two- or ten-days post-cFC. Motifs, associated transcription factors (TF), p-values and percentage of enrichment over background (bkg) are shown. (# of peaks for one day: 17,329, for two days: 11,288)
d) Bar chart of the percentage of TRAP+-enriched peaks containing one or less (violet) or more than one (blue) AP-1 motifs.

### Figure S5

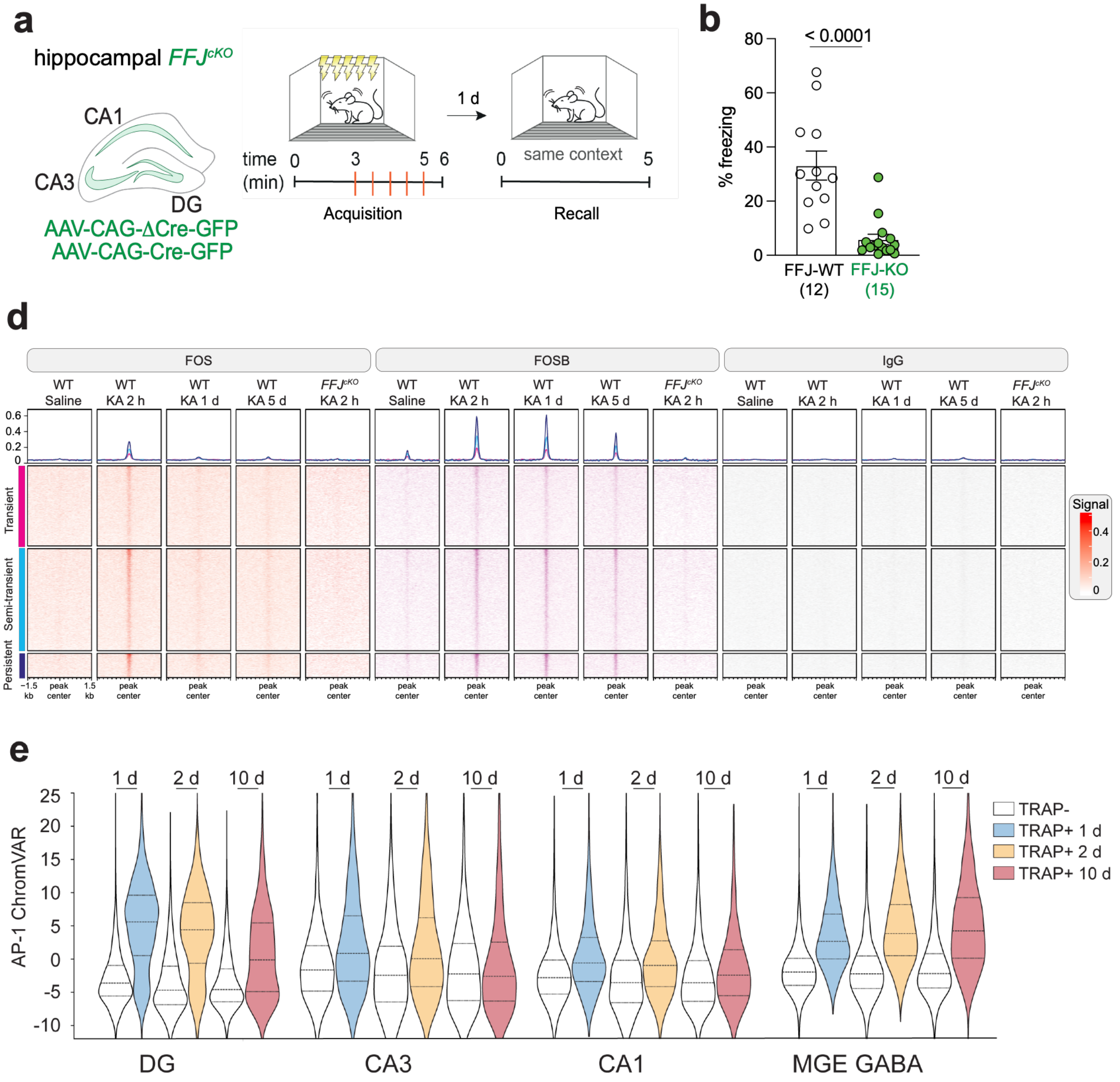


**Supplementary Figure 5: FOS family members as key players for ensemble neuron transcriptomic maintenance, related to Figure 3 and Figure 4**

a) Hippocampal knockout of *Fos*, *Fosb* and *Junb* (FFJ) via introduction of virally encoded Cre or control catalytically inactive Cre (ΔCre) recombinase and assessment of fear memory recall one day post-cFC.
b) Mean percentage of freezing upon five-minute memory recall ± SEM is displayed for FFJ-WT and FFJ-KO animals. (Numbers of individual animals are indicated, Mann-Whitney test for statistical comparison between conditions)
c) Aggregation plots and binding heatmap for CUT&RUN for FOS and FOSB from main Figure 3f, including IgG control.
d) Violin plots of AP-1 ChromVAR scores for all major cell classes (as in Fig. 4a).

### Figure S6

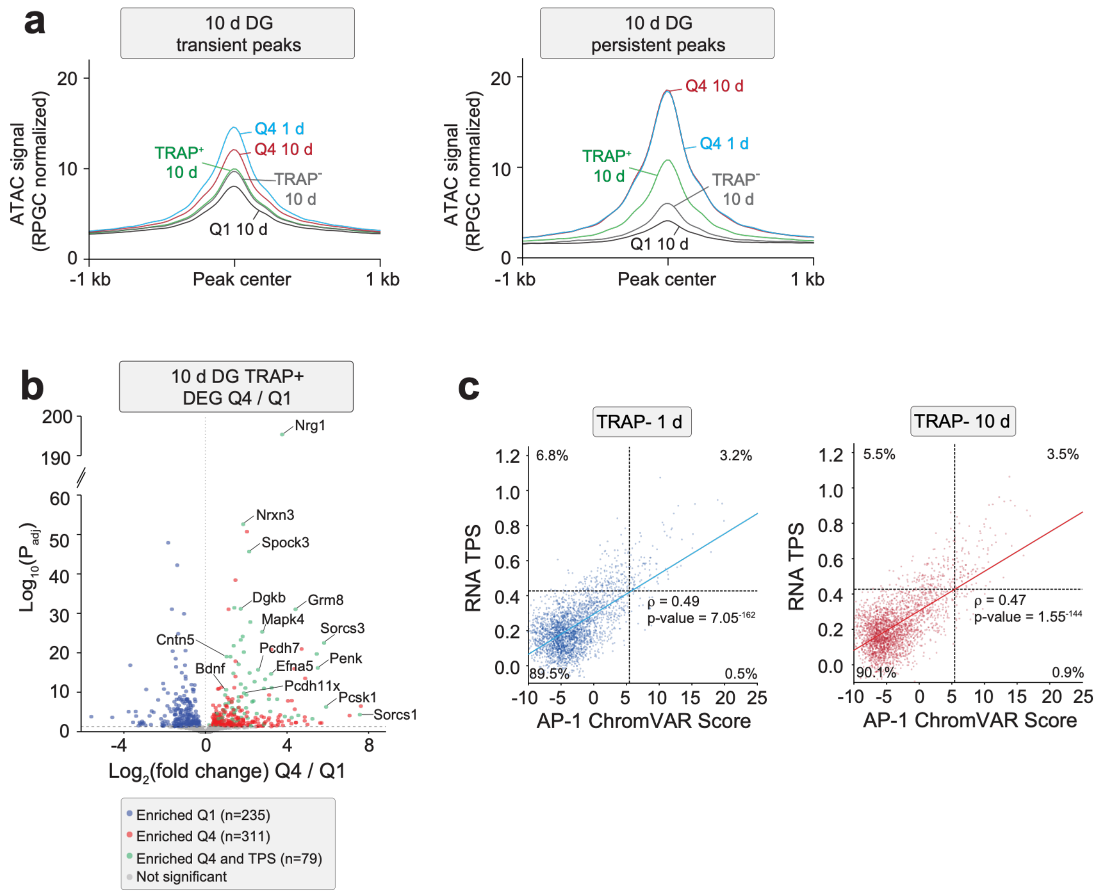


**Supplementary Figure 6: AP-1-associated regulatory elements promote ensemble-specific persistent gene expression, related to Figure 4**

a) Aggregate plot showing peak accessibility around the peak center for transient (left) or persistent (right) TRAP+-enriched peaks in the DG ten days post-cFC. Accessibility is displayed for either all TRAP+ neurons at ten days (green), or subsets of neurons, including AP-1 Low at ten days (black), AP-1 High (Q4) at one (blue) or ten (red) days.
b) Volcano plot of DESeq2 pseudobulked differential gene expression analysis between AP-1 High (Q4) and AP-1 Low (Q1) DG neurons ten days post-cFC. Blue and red dots indicate significantly down- or up-regulated genes, respectively (adjusted P-value <0.05, two-sided Wald test with Benjamini Hochberg correction for multiple testing). Genes that are enriched in Q4 and constitute components of the Transcriptional Program Score (TPS) are marked in green. Numbers of significant differentially expressed genes are indicated.
c) Correlation between TPS score and AP-1 ChromVAR score of TRAP- neurons at one (blue) or ten (red) days. Spearman's rank correlation coefficient (r), and percentage of neurons in each quadrant are indicated.

### Figure S7

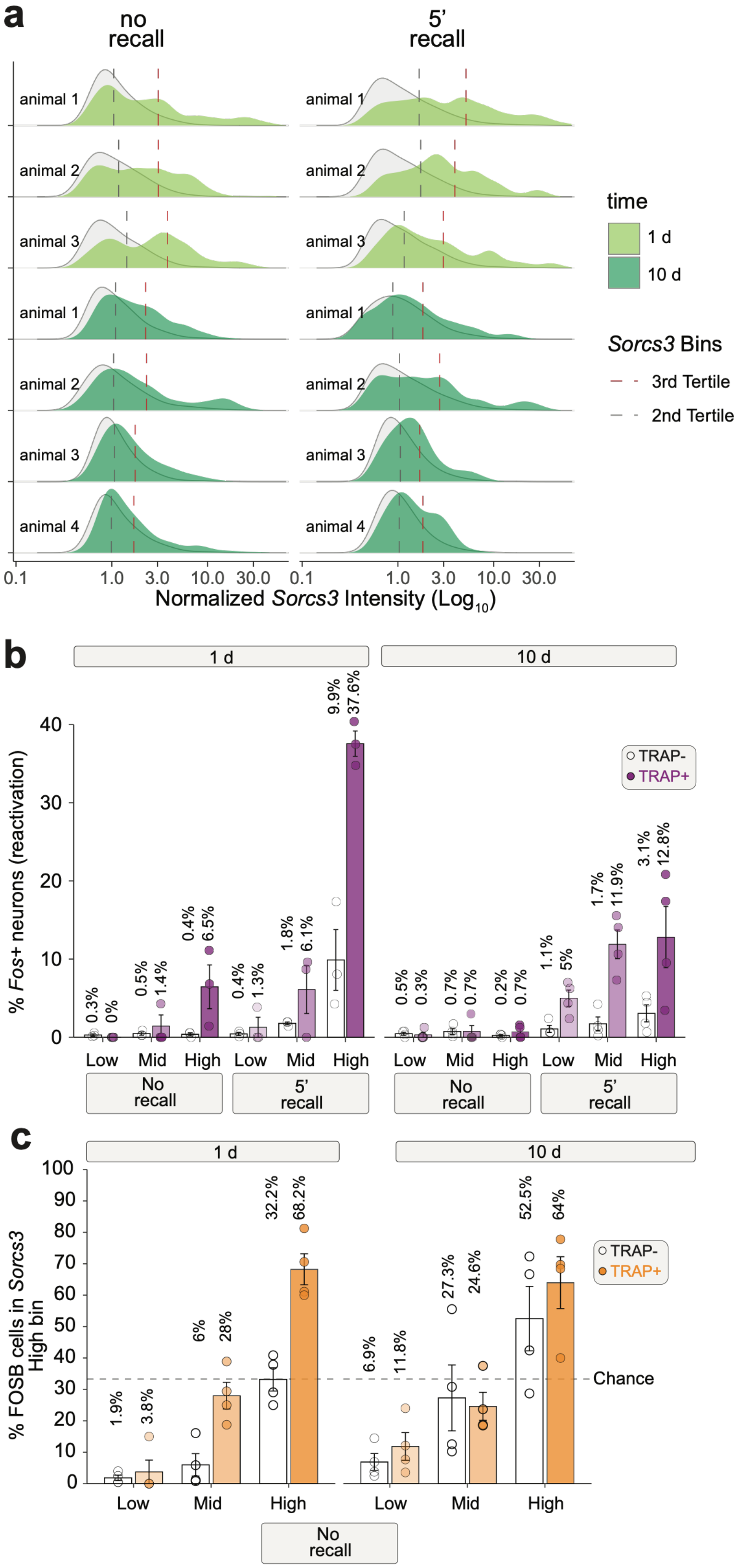
a
no recall
5' recall
animal 1
animal 2
animal 3
animal 1
animal 2
animal 3
animal 4
time
1 d
10 d
Sorcs3 Bins
3rd Tertile
2nd Tertile
0.1
1.0
3.0
10.0
30.0
Normalized Sorcs3 Intensity (Log10)
b
1 d
10 d
% Fos+ neurons (reactivation)
TRAP-
TRAP+
0.3%
0%
0.5%
1.4%
0.4%
6.5%
0.4%
1.3%
1.8%
6.1%
9.9%
37.6%
0.5%
0.3%
0.7%
0.7%
0.2%
0.7%
1.1%
5%
1.7%
11.9%
3.1%
12.8%
Low
Mid
High
No recall
5' recall
c
% FOSB cells in Sorcs3 High bin
1.9%
3.8%
6%
28%
32.2%
68.2%
6.9%
11.8%
27.3%
24.6%
52.5%
64%
Chance
No recall

**Supplementary Figure 7: *Sorcs3* mRNA levels correlate with rates of *Fos* reactivation and persistent FOSB expression, related to Figure 5**

a) Ridge density plots of $Log_{10}$ normalized *Sorcs3* FISH intensity values of individual animals analyzed. TRAP+ neurons are labeled in light (one day) or dark (ten days) green. TRAP- neurons from the same animal are depicted in light grey. Cutoffs for *Sorcs3* tertile bins used to assign neurons to *Sorcs3* low, mid or high are indicated with dashed lines.
b) Percentage of *Fos*+ neurons in TRAP+ and TRAP- cells that either fall within the *Sorcs3* high, mid or low tertile. One day no recall: N = 3 animals, TRAP+: 502 cells / TRAP-: 11444 cells; One day recall: N = 3 animals, n = TRAP+: 293 cells / TRAP-: 6180 cells; Ten day no recall: N = 4 animals, n = TRAP+: 757 cells / TRAP-: 10486 cells; Ten-day recall: N = 4 animals, n = TRAP+: 577 cells / TRAP-: 10529 cells.
c) Percentage of FOSB cells exhibiting low, mid or high FOSB protein expression in TRAP+ and TRAP- DG neurons that also display high *Sorcs3* mRNA expression. One day no recall: N = 4 animals, n = TRAP+ 215 cells / TRAP-: 2963 cells; Ten day no recall: N = 4 animals, n = TRAP+ 369 cells / TRAP-: 3196 cells.
